# Efficient inhibition of cell proliferation and promotion of apoptosis requires continuous treatment with abemaciclib

**DOI:** 10.1101/2021.11.15.468648

**Authors:** Torres-Guzmán Raquel, Ganado Maria Patricia, Mur Cecilia, Marugán Carlos, Baquero Carmen, Yang Yanzhu, Yi Zeng, Bian Huimin, Du Jian, de Dios Alfonso, Puig Oscar, Lallena Maria Jose

**Affiliations:** Discovery Chemistry Research & Technology, Eli Lilly and Company, Madrid, Spain; Eli Lilly and Company, Indianapolis, IN, USA; Eli Lilly and Company, New York, NY, USA

## Abstract

Abemaciclib is an oral, selective cyclin-dependent kinase 4 & 6 inhibitor (CDK4 & 6i), approved for hormone receptor-positive (HR+), human epidermal growth factor receptor 2-negative (HER2–) advanced breast cancer (ABC) as monotherapy for endocrine refractory disease, and with endocrine therapy (ET) for initial treatment and after progression on ET. Abemaciclib has also shown clinical activity in combination with ET in patients with high risk early BC (EBC). Here, we examined the preclinical attributes of abemaciclib and other CDK4 & 6i using biochemical and cell-based assays. In vitro, abemaciclib preferentially inhibited CDK4 kinase activity versus CDK6, resulting in inhibition of cell proliferation in a panel of BC cell lines with higher average potency than palbociclib or ribociclib. Abemaciclib showed activity regardless of *HER2* amplification and phosphatidylinositol 3-kinase (*PI3KCA)* gene mutation status. In human bone marrow progenitor cells, abemaciclib showed lower impact on myeloid maturation than other CDK4 & 6i when tested at unbound concentrations similar to those observed in clinical trials. Continuous abemaciclib treatment provided profound inhibition of cell proliferation, and triggered senescence and apoptosis. These preclinical results support the unique efficacy and safety profile of abemaciclib observed in clinical trials.

## Introduction

Breast cancer is the second most common cancer worldwide (1). Pharmacologically targeting cyclin-dependent kinase 4 and 6 (CDK4 & 6) has proven to be a successful therapeutic approach in patients with estrogen receptor-positive (ER+) breast cancer (2). D-type cyclins bind and activate CDK4 & 6. Once active, the holoenzyme phosphorylates the retinoblastoma tumor suppressor protein (Rb), causing the release of transcription factors that promote cell cycle progression to S phase, ultimately leading to cell proliferation (3). CDK4 & 6 are commonly dysregulated in cancer cells, promoting cell proliferation, and suppressing cell senescence (4, 5).

To date, the US Food and Drug Administration (FDA) has approved three CDK4 & 6 inhibitors (CDK4 & 6i) for treatment of hormone receptor-positive (HR+) metastatic breast cancer (MBC); palbociclib (PD0332991; Ibrance; 6), ribociclib (LEE011; Kisqali; 7) and abemaciclib (LY2835219; Verzenio; 8, 9). Moreover, abemaciclib is the first FDA-approved CDK4 & 6i approved for the adjuvant treatment of HR+, HER2–, node-positive, early breast cancer at high risk of recurrence and a Ki-67 score ≥20% (10, 11). Differences have been observed in both efficacy and severity of neutropenia among the available CDK4 & 6i, generating interest in a possible mechanistic explanation (12).

Abemaciclib is an ATP-competitive, reversible, selective CDK4 & 6i, which when administered as monotherapy, had a safety profile enabling continuous dosing in a Phase 1 clinical study (NCT01394016) (13). The findings were observed in previously treated patients with HR+ MBC, non-small cell lung cancer, and melanoma (14). Moreover, antitumor activity of abemaciclib as a single agent in HR+, HER2– MBC has been demonstrated in a Phase 2 trial (NCT02102490) (15). The efficacy and safety of combining abemaciclib plus endocrine therapy (ET) in HR+ MBC has also been demonstrated in key Phase 3 trials, MONARCH 2 and MONARCH 3 (16, 17). Abemaciclib is the only CDK4 & 6i to demonstrate significant invasive disease-free survival improvement in the adjuvant treatment of patients with high risk, early breast cancer (EBC) when administered with standard ET (18).

Based on previously reported differences between CDK4 & 6i, their efficacy, and their impact on neutropenia, this study examined the preclinical biochemical and cellular profiles of abemaciclib, palbociclib, and ribociclib in a panel of BC cell lines, their impact on neutrophil maturation and the results of intermittent versus continuous treatment. Our results demonstrate that abemaciclib has unique pharmacological properties that are consistent with the safety/efficacy profile observed in clinical trials.

## Materials & Methods

### Ethics statement

Samples included in this study were provided by the Biobank Hospital Universitario Puerta de Hierro Majadahonda (HUPHM)/Instituto de Investigación Sanitaria Puerta de Hierro-Segovia de Arana (IDIPHISA) (PT17/0015/0020 in the Spanish National Biobanks Network). Samples were processed following standard operating procedures with the appropriate approval of the Ethics and Scientific Committees.

### Human blood samples

Human whole blood was obtained from six healthy donors who previously provided written informed consent. Mature neutrophils were isolated from human whole blood, using a negative-selection technique (MACSxpress isolation kit, Miltenyi 130-104-434). CD16 cell surface expression (Miltenyi, 130-113-396) was used as quality control; only samples with yield > 95% passed QC. CD34+ human bone marrow primary progenitor cells were obtained from Tebu bio (BM34C-4).

### Materials

Unless otherwise indicated, all preclinical data described herein were obtained using the methanesulfonate salt of each compound for abemaciclib and palbociclib, and the succinate salt for ribociclib. Full chemical names have been previously published (19). See Supplementary Materials for full list of materials.

### Biochemical characterization

#### TR-FRET assays

Compounds were mixed with kinase solution at 37.5X final assay concentration and 1% dimethyl sulfoxide (DMSO) and incubated for 30min. Then, the reactions were rapidly diluted in a saturating concentration of ATP and the TR-FRET signal was measured continuously in a multiplate reader Envision (Perkin Elmer).

The signal of the baseline control was subtracted from the maximum signal (without compound), as well as from each test compound value. This corrected emission ratio (ER^*^) was used to calculate the percentage of activity recovery for each condition:

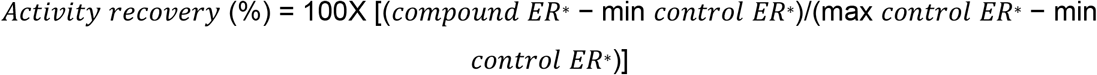

Ki values were calculated using the following equation: *K*_*i*_ = *IC*_50_ /[1 + (*S* /*K*_*m*_)] To measure IC_50_ values using TR-FRET, a LANCE® Ultra kinase assay was used. 5 μL of test compound was mixed with 5 μL of kinase and 5 μL substrate-antibody mixture. Sample containing the peptide, ATP, antibody, and kinase without inhibitor were used as the high reaction control and wells containing the peptide, ATP, and antibody were used as the low reaction control. Test compounds were pre-incubated with the kinase for 30min. The substrate-antibody solution was then added and incubated 1h at room temperature (RT). Test compounds were run in duplicate. Assay was read in a multiplate reader Envision (Perkin Elmer). Results were fitted to a dose-response curve with variable slope and constraints of 0 and 100 for bottom and top, respectively, to generate the IC_50_ values.

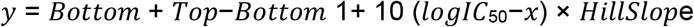

#### Filter binding (FB) assays

Compounds were mixed with substrate mix (C-terminal retinoblastoma fragment [CTRF] peptide and ATP/33-P ATP) at a final concentration range of 2 μM-0.1 nM. The mix was incubated for 90min at RT and the reaction was then stopped by adding 80 μl of 10% ortho-Phosphoric acid. The mix was next transferred to Multiscreen filter plates (Millipore) to retain phosphorylated peptide. The plates were read in a Microbeta Trilux instrument. Reaction mix with excess of EDTA was used as assay “bottom signal”; complete reaction mix without the inhibitor/compound was used as assay “top signal”.

For IC_50_ determination, results were fitted to a four-parameter dose-response curve using the following equation:

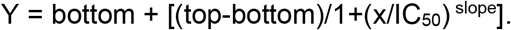

For the ATP saturation studies K_i_ and K’_i_ parameters were estimated using the formula:

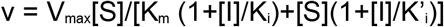

### Cell lines and culture condition

A panel of 37 BC cell lines (see Table S3 for details) was obtained from the American Type Culture Collection (ATCC, Manassas, Virginia, USA 30-4500 K) or the German Collection of Microorganisms and Cell Cultures GmbH (DSMZ). Cells were cultured according to ATCC or DSMZ recommendations for fewer than ten passages. Cells were seeded in 96 or 384-well plates and incubated overnight prior to treatment.

CD34+ cells (Figure 3, S4-6) were thawed and resuspended in IMDM, 10% hiFBS (Fisher), and 1% Pen/Strep (Invitrogen), supplemented with GMCSF 10 ng/ml, G-CSF 10 ng/ml, SCF 100 ng/ml, IL3 10 ng/ml and IL6 10 ng/ml.

Mature neutrophils were plated in a 96-well plate (60K cells per well) in 100 μl RPMI 1640 (Invitrogen), 10% hiFBS (Fisher) and 1% Pen/Strep (Invitrogen).

### In vitro drug treatment

#### Breast Cancer Cell panel

Cells were plated at 1,000 or 2,000 cells per well in 384-well plates in a total volume of 25 μl of either growth media alone; with 2 μM staurosporine; or with decreasing concentrations of test compounds with a range 2 μM-0.1 nM. Test compounds were prepared in 100% DMSO using a dilution factor of 1:3 before addition to the cells pre-diluted in growth media. Cells were incubated at 37°C until the untreated cells had doubled twice. Cells were fixed with 70% ethanol, treated with RNAse and nuclei stained with Propidium Iodide (PI). Cell nuclei per well were counted using an ACUMEN EXPLORER™ (STP LABTECH LTD) to determine the number of cells remaining after treatment.

#### Mature neutrophils from blood

Neutrophils were plated in a 96-well plate (60K cells/well) in the above-mentioned cell medium. A 10 mM stock of each CDK4 & 6i was used to make ten-point 1:3 serial dilutions in 100% DMSO; multiple dilution steps were used to reach the concentration in the assay: 10 μM, 0.1% DMSO. Non-treated cells (0.1% DMSO) and roscovitine (20 μM) were used as controls. Neutrophils were incubated for 6 h. Following treatment, cells were washed with FACS flow (BD, 342 003), and supernatant was removed. The cell pellet was stained with Annexin V-FITC antibody for 10 min in the dark. Cells were washed with Annexin buffer, centrifuged at 300 G for 5min, supernatant was discarded, and cell pellet was resuspended in 100 μl of Annexin buffer and analyzed by flow cytometry. PI 1:200 was added automatically by the cytometer. All incubations occurred at 37ºC, 5%CO_2_.

The percentage of alive, early apoptosis, late apoptosis, and dead cells was monitored using the Annexin V/PI assay. Cells were incubated with an anti-human Annexin V-FITC antibody, binding specifically to phosphotidylserine (PS), for 10 min in the dark. PS redistributes to the outer part of the cell membrane in an early stage of apoptosis; in late apoptotic cells, as the cell membrane loses its integrity, both PS and PI can be detected. Thus, the four different phases of apoptosis can be distinguished using flow cytometry technology: Annexin V-/PI- corresponds to alive cells, Annexin V+/PI- corresponds to early apoptotic cells, Annexin V+/PI+ corresponds to late apoptotic cells and Annexin V-/PI+ corresponds to the dead subpopulation of cells. The total number of cells (early, late and dead cells) are represented.

#### CD34+ bone marrow progenitor cells

CD34+ progenitor cells were exposed to abemaciclib, palbociclib or ribocilib and the amount of cells/ml were measured after 13 days. On Day 0, 5 000 CD34+ progenitor cells/well were seeded in a deep well plate and were incubated overnight. On Day 1, cell medium was changed for IMDM, 10% hiFBS (Fisher) and 1% Pen/Strep (Invitrogen), supplemented with GMCSF 10 ng/ml, G-CSF 10 ng/ml, SCF 100 ng/ml, IL3 10 ng/ml and IL6 10 ng/ml. Cells were treated with DMSO (vehicle) or with 3-fold dilutions of compounds (abemaciclib, palbociclib, or ribociclib) at a range of concentrations of 20 μM-1 nM and incubated for 13 days. On Days 3, 6 and 10 the cell medium and compounds were renewed (Days 3&6) or supplemented (Day 10) to maintain an optimal cell density. To compare the effects of each individual compound at the Cmax, fraction unbound, results at 26 nM (abemaciclib and all active metabolites), 38 nM (palbociclib) or 1 548 nM (ribociclib) were interpolated (14, 20-23).

#### T47D cells: Continuous dosing

T47D cells were seeded in six-well plates (50 000 cells/well) in 2 ml of cell medium (10% hiFBS [Fisher] and 1% Pen/Strep [Invitrogen]). On Day 1, cells were dosed with abemaciclib or palbociclib at 250 nM, 100 nM or 50 nM final concentrations. Non-treated cells (0.04% DMSO) and staurosporine 1 μM were used as controls for minimum and maximum inhibition controls respectively. After 2, 6, or 9 days of incubation, cell medium in supernatant was collected and transferred to a deep well plate. Cells were then detached using trypsin and transferred to the deep well plate and spun at 1 300 rpm for 5 min. Cell pellet was washed using PBS1X and plated for apoptosis measurement (Annexin V/PI), using the procedure described above (mature neutrophils). All cells were gated on FSC/SSC to exclude debris and doublets.

#### T47D cells: Washout study

100 000 T47D cells/well (short treatment) or 15 000 T47D cells/well (long treatment) were seeded in six-well plates (Thermo Fisher, 140675) in 2 ml of cell medium (10% hiFBS [Fisher] and 1% Pen/Strep [Invitrogen]). The plates were incubated overnight. On Day 1 cells were dosed with 100 nM or 50 nM of abemaciclib or palbociclib as a single treatment or in combination with 5 nM of 4OH-tamoxifen in triplicates. Non-treated cells (DMSO 0.1% as vehicle), as well as staurosporine (100 nM) or Mytomycin C (200 nM), were used as controls. After 2 or 8 days of continuous treatment the compounds were removed, and cells were incubated with cell medium for four more days. On Days 2 or 8 of continuous treatment, or 4 days after compound removal, cells were detached using Accutase for 10 min. All supernatants were collected during the detachment steps to prevent the loss of cells in the suspension. Cells were transferred to a deep well plate and divided into four aliquots for analysis. Cell sensitivity, apoptosis, and senescence were measured by number of cells (remaining after treatment), Annexin V/PI, or cell even green assays respectively. Mitosox was also measured (Sup. Mat.).

For fluorescent detection of β-galactosidase, cells were washed with PBS 1X and fixed with 2% PFA (Acros organics, 119690010) for 10 minutes. Cells were then washed with PBS+1%BSA and incubated with the cell even green reagent (Thermo, C10841) following the vendors indications (2h, 31ºC, no CO_2_). Finally, cells were washed and resuspended in 1% BSA in PBS for FACS analysis (488-nm laser and 525/50 nm filter). For cell proliferation inhibition (cell number), data was normalized versus non-treated cells and staurosporine-treated cells and the percentages were plotted with Graph Pad v8.4.3. Senescence (cell even green) was represented as the percentage of green positive cells and compared to non-treated cells.

### Western blot

Cells were washed with PBS and lysed with ice-cold lysis buffer containing 2 mM PMSF, 10 mM EDTA, and 2xHalt protease and phosphatase inhibitor cocktail (Thermo). After a 10 min incubation on ice, the lysate was centrifuged at 14 000 rpm at 4°C for 5 min. The supernatant was stored in –80°C for Western blotting. 45μg of each sample quantified by BCA assay was loaded onto NuPAGE 4% to 12% Bis-Tris Gel and immobilized onto nitrocellulose membrane using Tris-Glycine transfer buffer at 100 V for 1 h. The immunoblotting was performed in a blocking buffer of 2.5% non-fat milk/TBS-T and detected by anti-CDK4 antibody and anti-CDK6 antibody using ECL-HRP on Fujifilm LAS4000.

### Analysis and statistical considerations

Raw data were analyzed with FlowJo 10.6 software. Graph Pad v8.4.3 software was used for data analysis and representation of final readouts. JMP (Statistical Discovery from SAS) was used for the statistical treatment (ANOVA, pairwise analysis) of the data.

The IC_50_ was determined by curve fitting to a four-parameter logistic equation for each output using GENEDATA SCREENER® tool: or GraphPad Prims®.

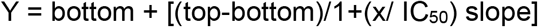

where Y = % inhibition (%Inh = [(median Max- x/ median Max – median Min)]/100], X = concentration yielding y% inhibition.

Ten-points curves were obtained and a RelIC_50_ was calculated.

### Flow cytometry analysis

Cells were analyzed using a flow cytometer (Macsquant Analyzer 10, Miltenyi). Events were gated for debris exclusion and singlets selection. A minimum of 3 500 cells were analyzed per sample. PI (1:200) was added automatically by the cytometer. Cells negative for Annexin V and PI (% alive cells) were used for analysis. Cells/ml (gated on singlets) were evaluated using the Macsquantify 2.13 software (more details on analyses in Sup. Mat.).

## Results

### 1. Abemaciclib is a more potent inhibitor of CDK4 than CDK6

The potency of abemaciclib, palbociclib and ribociclib to inhibit CDK4/cyclinD1 and CDK6/cyclinD3 activity was evaluated by measuring retinoblastoma phosphorylation by TR-FRET or FB assays. Abemaciclib is a highly potent inhibitor of CDK4/cyclin D1 showing in TR-FRET assays an IC_50_ value of 0.94 nM; abemaciclib potency against CDK6/cyclinD3 was 14 nM, demonstrating a selectivity ratio CDK4/CDK6 of 15-fold. In contrast, palbociclib yielded a selectivity ratio CDK4/CDK6 of 0.3-fold and ribociclib 4.3-fold (Figure 1A). These results were consistent regardless of the method used (FB assays IC_50_ or Ki determination by FB or TR-FRET; Table S1).

**Figure 1.**
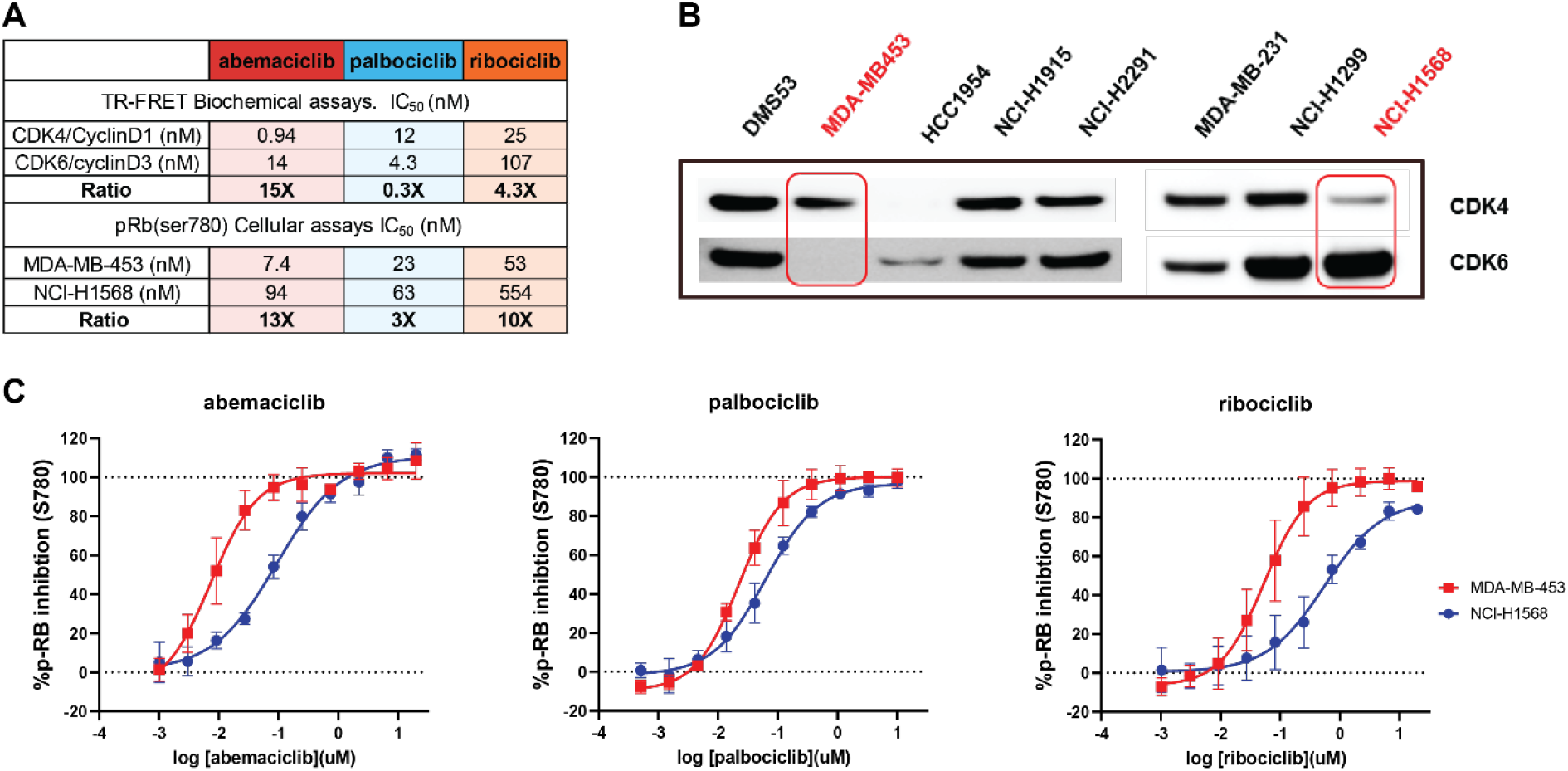
Abemaciclib is a more potent inhibitor of CDK4 than CDK6 in both biochemical and in vitro cell-based assays. In a head-to-head comparison, abemaciclib showed higher potency to inhibit CDK4 than CDK6 with a broader margin of selectivity than other CDK4 and CDK6 inhibitors. This was confirmed in biochemical assays and in cellular systems. Biochemical potency (KiATP) and selectivity ratio CDK6/CDK4 (additional IC_50_ values, FB assay and TR-FRET in Supplemental Material) as cell assays for abemaciclib and palbociclib are included in **table A)**. Figure **1B**) represents the Western Blot showing the expression of CDK4 and CDK6 in different cell models. MDA-MB-453 and NCI-H1568 cells (highlighted) showed preferential expression of CDK4 or CDK6 respectively. **(C)** Dose-response curves of abemaciclib, palbociclib, and ribociclib showing inhibition of Rb (retinoblastoma) ser780 phosphorylation, quantitated intracellularly by high content imaging in CDK4 and CDK6 dependent cell models (MDA-MB-453 and NCI-H1568) Data reported as an average of four independent determinations (n = 4) ± standard deviation (SD; error bars). TR-FRET: Time-Resolved Foerster Resonance Energy Transfer; FB: Filter Binding.

**Figure 2.**
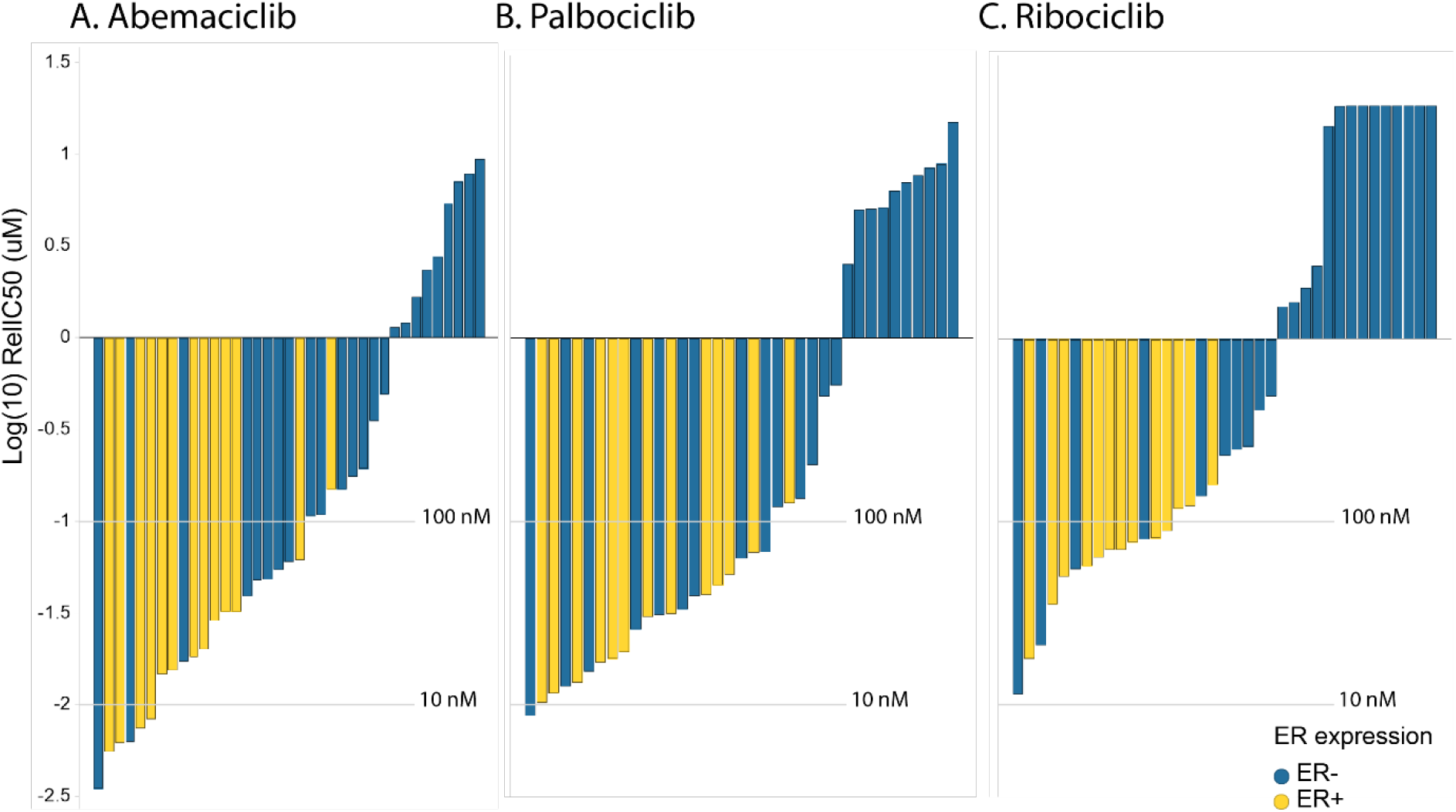
Abemaciclib shows greater potency than palbociclib & ribociclib in breast cancer cells. In vitro drug response waterfall plots for **(A)** abemaciclib, **(B)** palbociclib, **(C)** or ribociclib in a panel composed of 40 cell lines, either ER- (blue) or ER+ (yellow). Bar graph of log IC_50_ values (uM) and cell type. Cell lines are color coded by subtype: yellow is luminal ER+; blue is ER-. Waterfall plots were generated using the geometric mean for each cell line and treatment.

**Figure 3.**
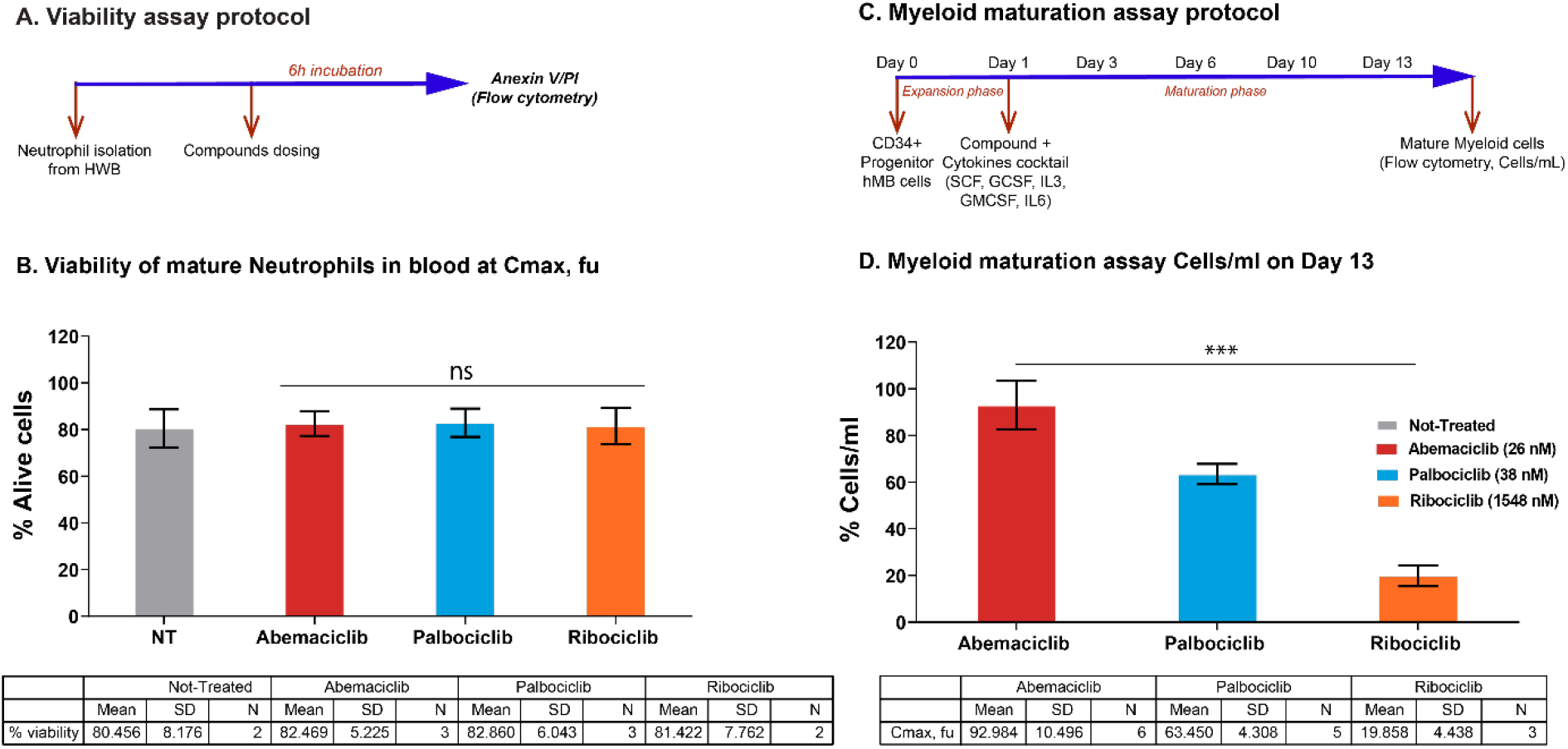
Impact on neutrophils maturation is lower upon abemaciclib treatment comparing with others CDK4 & 6 inhibitors in preclinical models. **(A-B)** Viability of isolated mature neutrophils from human whole blood at Cmax, fu. **(C-D)** Myeloid maturation assay, measuring cells per mL on Day 13. Data are plotted as the mean +/-SD of more than two independent replicates. HWB: Human Whole Blood, hBM: human Bone Marrow, PI: Propidium Iodide, Cmax,fu: maximum unbound concentration in plasma. *** p-value < 0.0001; ns. or no *: non-significant. One way ANOVA among three groups. In a pairwise comparison there is a size effect of 18.54 for abemaciclib versus palbociclib and 61.99 for abemaciclib versus ribociclib with p-values of 0.0035 and < 0.0001 respectively.

These results were confirmed in an isoform-specific cellular context using cell models that are dependent on CDK4 or CDK6, according to the relative expression of each kinase (Figure 1B). Intracellular phosphorylation of Rb in its residue 780 (Ser780) was measured using fluorescently labeled phospho-specific antibodies and high content imaging. Abemaciclib showed a dose-response inhibition of the Rb phosphorylation in the CDK4-dependent cell line MDA-MB-453 (Figure 1B) with a potency of 7.4 nM, and in the CDK6 dependent cell line NCI-H1568 with a potency of 94 nM (Figure1A-C), which translates to a selectivity ratio CDK4/CDK6 of 13-fold. In contrast, in same cellular environment, palbociclib and ribociclib showed lower potency 23 nM and 53 nM with a selectivity ratio of 3-fold and 10-fold, respectively. (Figure1C). CDK4 dependency in this cell line was confirmed by knocking out CDK4 or CDK6 by shRNA (Figure S2A). The higher potency shown by abemaciclib in inhibiting Rb phosphorylation was reproducible across several breast cancer cell lines (Table S2).

Additionally, to understand whether the profound inhibition of CDK4 may results in a more durable response, the effects of abemaciclib and palbociclib on Rb phosphorylation were assessed in an in vitro washout experiment (Figure S1A). In the breast cancer cell line MDA-MB-453, intracellular CDK4 activity was functionally inhibited during the treatment phase, as observed by the complete depletion of pRb (Ser780) regardless of treatment (Figure S1B). Under these conditions, both compounds showed relative IC_50_ values for pRb (Ser780) reduction in the low nM range: 6.4 nM for abemaciclib and 14.1 nM for palbociclib (Figure S1C). After compound removal, Rb phosphorylation was recovered in cells previously exposed to palbociclib, but not in those treated with abemaciclib, which maintained a relative IC50 below 100 nM even 12 h after compound removal. In cells pre-treated with palbociclib, relative IC_50_ increased to 446.7 nM 12 h after compound removal (Figure S1B-C). Taken together, those results demonstrate the sustained target inhibition and longer-term effect of abemaciclib in ER+ BC cell lines.

### 2. Abemaciclib is a potent inhibitor of proliferation in breast cancer cell lines

The impact of the three CDK4 & 6i on both mature neutrophils and myeloid maturation was investigated in vitro to gain insights on the effects in hematopoietic cells at a concentration similar to the plasma levels of these inhibitors in clinical trials. For abemaciclib, we combined the parent molecule with its two active circulating metabolites with equivalent pharmacology (14, 20-23). At the concentration (corrected for protein binding) equivalent to the average clinical Cmax reported for each CDK4 &6i (unbound Cmax), there was no impact on circulating neutrophils in blood (Figures 3A-B). However, the impact on the maturation of progenitor cells from bone marrow was lower with abemaciclib treatment compared with palbociclib or ribociclib (Figures 3C-D). Furthermore, in human bone marrow progenitor cells, abemaciclib triggered apoptosis with lesser extent than palbociclib (Figure S4) and had also a lesser impact on neutrophil maturation markers (Figure S5). Importantly, neither known abemaciclib metabolites nor combination of abemaciclib with fulvestrant had any impact on neutrophils maturation in vitro (Figures S5 and S6).

### 3. Abemaciclib showed a lesser impact on myeloid maturation than other CDK4 & 6 inhibitors

The impact of the three CDK4 & 6i on both mature neutrophils and myeloid maturation was investigated in vitro. At the Cmax fraction unbound for each CDK4 & 6i (14, 20-23), there was no impact on circulating neutrophils in blood (Figures 3A-B). However, the impact on the maturation of progenitor cells from bone marrow was lower with abemaciclib treatment compared with palbociclib or ribociclib (Figures 3C-D). Furthermore, in human bone marrow progenitor cells, abemaciclib triggered apoptosis with lesser extent than palbociclib (Figure S4) and had also a lesser impact on neutrophil maturation markers (Figure S5). Importantly, neither known abemaciclib metabolites nor combination of abemaciclib with fulvestrant had any impact on neutrophils maturation in vitro (Figures S5 and S6).

### 4. Prolonged treatment with abemaciclib leads to apoptosis

To understand the impact of longer treatment of abemaciclib and palbociclib on their efficacy in breast cancer cell lines, a time course (2, 6 and 9 days) at 50, 100 and 250 nM was performed to evaluate the biological effect by flow cytometry, early (PI-/Annexin V+) and late apoptosis (PI+/Annexin V+), and cell death (PI+/Annexin V-) events (Figure 4). Early apoptotic effects are observed at Days 2 and 6, and late apoptotic effects are significant upon 9 days of treatment (Figure 4). Although the increase of apoptosis is observed upon both CDK4 and CDK6 inhibitors after 8 days of continuous dosing, the percentage in total apoptotic and dead cells is higher upon abemaciclib treatment compared with palbociclib at the two concentrations tested (28.1% versus 13.2% at 50 nM; 40.7% versus 18.7% at 100 nM, for abemaciclib and palbociclib, respectively). Taken all those data together we conclude that prolonged treatment with abemaciclib promoted dose-dependent apoptosis in T47D (Figure 4) and MCF7 breast cancer cells (data not shown).

**Figure 4.**
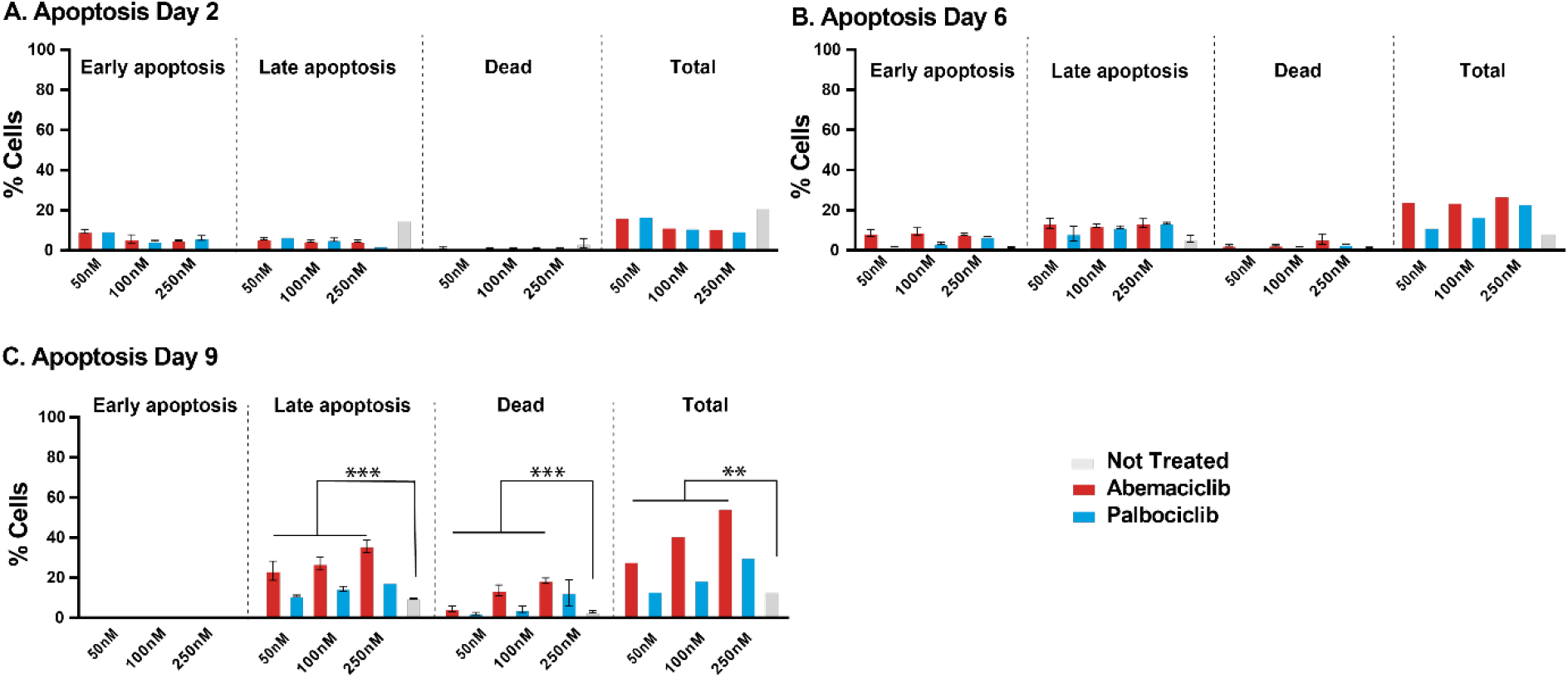
Prolonged treatment of breast cancer cells with a CDK4 & 6 inhibitor is necessary to sustain cell growth inhibition and promote apoptosis. T47D cells were treated with DMSO, 50, 100 or 250 nM of the CDK4 & 6i abemaciclib or palbociclib for 2, 6, and 9 days. **(A-C)** percentage of apoptotic cells are monitored by Annexin V and PI. The data are plotted as the mean (+/- SD) of three experiments for CDK4 & 6i treatment, and the mean of three experiments (+/- SD) for untreated samples. ANOVA analysis of Day 6 data show that % cell in apoptosis (late, total, or dead) is significantly different between groups treated with abemaciclib or not treated (FC 2.9, 3.1 and 3.3 respectively); % cell in late apoptosis is significantly different between groups treated with palbociclib or not treated (FC 1.45), although there is not significant change in the case of total apoptosis or dead cells in groups treated with palbociclib or not treated (FC 1.51 and 1.48 respectively). ANOVA: Analysis of Variance, SD: Standard deviation, FC: Fold Change. *** p-value ≤ 0.0001, ** p-value ≤ 0.001; if no p-value presented, the differences between groups were not statistically different.

### 5. Permanent exposure leads to durable effects after compound removal

To compare the persistence of effects after compound removal, T47D cells (ER+, PR+, HER2–) were treated with a CDK4 & 6i (abemaciclib or palbociclib) as monotherapy or in combination with tamoxifen for 2 to 8 days (Figure 5A). Compounds were then removed, and cells were incubated for 4 more days. Cell proliferation, senescence and apoptosis were monitored (Figure 5 and Figures S7&S8). Overall, the effect was more durable when cells had previously been treated for longer (8 days). Nevertheless, it is worth highlighting that after 2-day treatment with abemaciclib plus tamoxifen, a higher cell proliferation inhibition was observed (Figure 5B), which is consistent with previous data obtained in monotherapy (Figure S8A).

**Figure 5.**
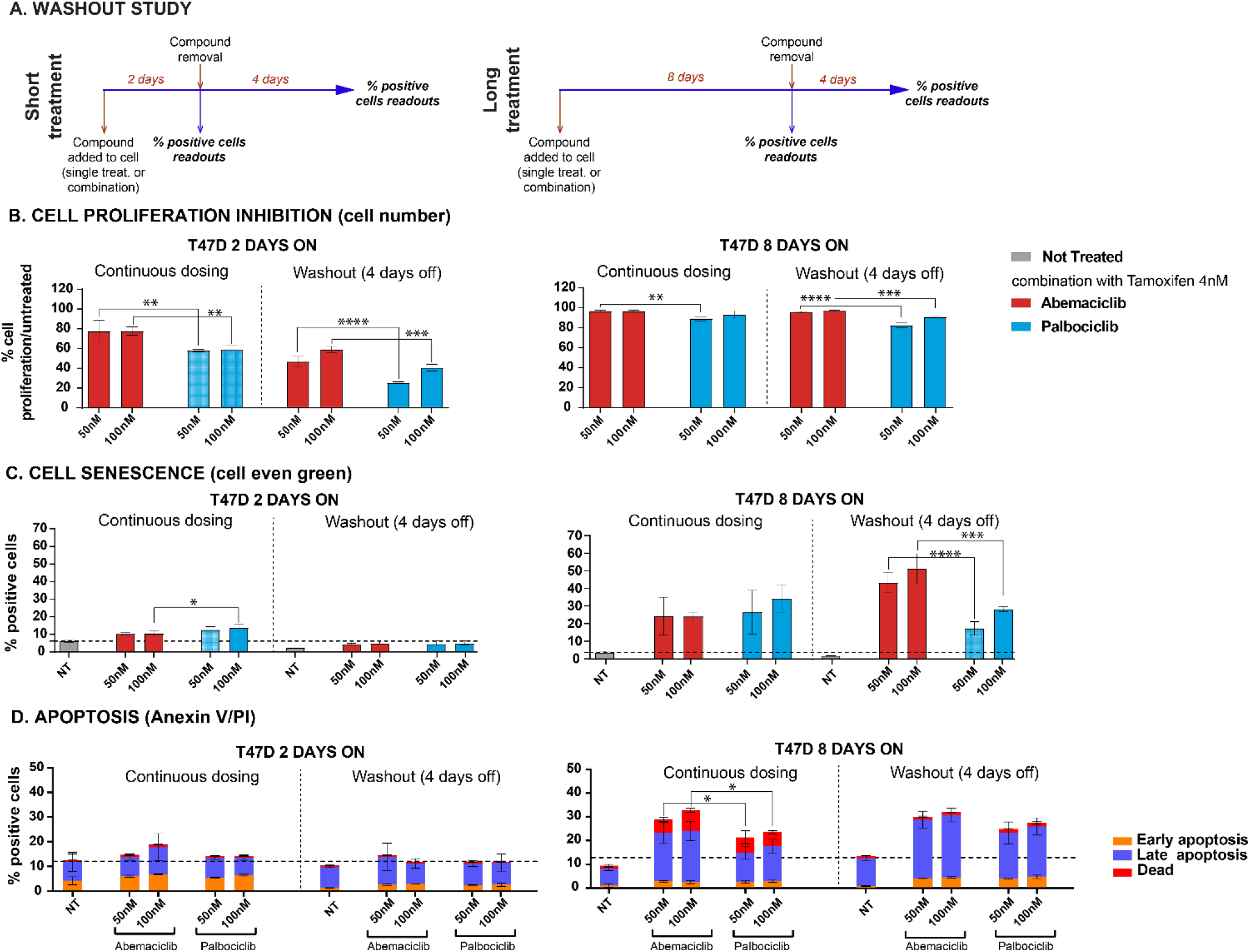
Washout studies demonstrated durable effects after abemaciclib removal. Cells treated with either abemaciclib or palbociclib + 5 nM tamoxifen. **(A)** Cell proliferation inhibition. **(B)** Percentages of senescent cells were monitored by cell even green kit. **(C)** Percentage of apoptotic cells, monitored by Annexin V and PI. The data are plotted as the mean (+/- SD) of three experiments. NT: non-treated. * p-value ≤ 0.05; ** p-value ≤ 0.01; *** p-value ≤ 0.001; **** p-value ≤ 0.0001; ns. or no *: non-significant.

When comparing the different treatments, no significant difference in cell number inhibition after compound removal (8-day treatment + 4-day WO) was observed with tamoxifen plus either abemaciclib or palbociclib (Figure 5B). However, the number of senescent cells remained significantly higher with the combination of abemaciclib + tamoxifen compared to combinations with palbociclib (Figure 5C). A significant number of apoptotic cells remained after compound removal (8-day treatment + 4-day WO; Figure 5D). These results were consistent regardless of the methodology used (cell even green or Mitosox) and similar results were observed with monotherapy treatments (Figure S8) or in combination with fulvestrant (data not shown).

## Discussion

CDK4 & 6i have demonstrated robust clinical activity, receiving FDA approval for the treatment of advanced BC (11, 24, 25), and are being further investigated in additional clinical trials. Here, we examined the preclinical biochemical and cellular profiles of abemaciclib, palbociclib, and ribocilib in a panel of BC cell lines, neutrophils, and bone marrow progenitor cells. We studied the activity profile of each agent in broad panels of BC cells in single treatment, as well as in different dose regimen schedules.

In cell-free assays, abemaciclib showed selectivity for CDK4 over CDK6; in cell-based assays, it preferentially inhibited the proliferation of cells dependent on the presence of CDK4, not CDK6, and it showed to be a more potent CDK4 inhibitor and selective against CDK6 than palbociclib and ribociclib (26)(Figure 1, Table S1). The biochemical profile translated to cell-based assays, with CDK4-dependent breast cancer cell lines showing a profound inhibition of pRb under abemaciclib treatment. Robust cell cycle arrest, through inhibition of pRb, correlated with potent inhibition of cell proliferation in a large panel of BC cell lines. For instance, abemaciclib demonstrated greater potency than palbociclib and ribociclib in BC cell lines, regardless of ER positivity. Additionally, abemaciclib potently inhibited cell proliferation in BC cell lines, regardless of *HER2* amplification or *PI3KCA* and *BRCA1/2* gene mutation status.

Clinically, lower incidence and severity of neutropenia was reported in patients receiving abemaciclib treatment, compared to patients receiving either palbociclib or ribociclib (14, 27). The relationship of CDK6 and cyclinD3 in the maturation of the myeloid cells has been broadly discussed and demonstrated in preclinical studies with transgenic mice models lacking either CDK4 or CDK6 in adult hematopoiesis (28-31). In vitro, we investigated the impact of the three CDK4 & 6i on both mature neutrophils and the myeloid maturation. None of the CDK4 & 6i impacted mature circulating neutrophils in the bloodstream; however, abemaciclib treatment resulted in a lower impact on the maturation process compared to palbociclib and ribociclib, which may contribute to lower incidence of neutropenia. The higher and more selective activity of abemaciclib against CDK4 than CDK6 (Figure 1, Table S1), as well as the ratio of unbound Cmax to CDK6 potency, may explain the different rates of neutropenia observed among treatments (14). The differentiated pharmacological profile of abemaciclib may contribute to its tolerability profile that allows for continuous dosing whereas palbociclib and ribociclib need to be administered intermittently.

In vitro, treatment with abemaciclib and palbociclib inhibited cell proliferation by inhibiting Rb phosphorylation. After compound removal, Rb phosphorylation inhibition was maintained only in cells treated with abemaciclib, demonstrating the sustained target inhibition and longer-term effect of abemaciclib in ER+ BC cell lines. Consistent with this observation, T47D cells treated with abemaciclib were inhibited for longer, after compound removal, than cells treated with palbociclib. The anti-proliferative activity of abemaciclib thus had a more durable effect in promoting apoptosis, emphasizing the importance of efficient target inhibition in leading to a durable cellular response. Continuous treatment with abemaciclib promoted a greater response than intermittent treatment, as monitored by remaining cell number, senescence, and cell apoptosis. This suggests that continuous treatment is required to observe complete senescence and irreversible effects through apoptosis. We conclude that in preclinical models, continuous treatment with abemaciclib is required for profound and sustained effects, resulting in superior activity.

## Conclusion

In preclinical experiments, abemaciclib is a potent, selective cell growth inhibitor, inhibiting preferentially the CDK4/Cyclin D1 complex and leading to cell senescence and cell death in breast cancer cell lines with broad molecular profiles. Abemaciclib has a lesser impact on neutrophils maturation in vitro than other CDK4 & 6, which is consistent with lower incidences of neutropenia observed in clinical settings and may allow for a prolonged treatment. After prolonged dosing with abemaciclib, cells show sustained inhibition of cell proliferation that leads to irreversible effects through apoptosis. These preclinical results support the differentiated safety and efficacy profile of abemaciclib observed in clinical trials.

## Table of Abbreviations

ATCC: American Type Culture Collection
BC: Breast cancer
BCA: Bicinchoninic acid
CTRF: C-terminal retinoblastoma fragment
DMSO: Dimethyl sulfoxide
DSMZ: German Collection of Microorganisms and Cell Cultures GmbH
EDTA: Ethylenediaminetetraacetic acid
ER: Estrogen receptor
ET: Endocrine therapy
FB: Filter Binding
FDA: Food and Drug Administration
hBM: Human bone marrow
HR: Hormone receptor
HWB: Human whole blood
IMDM: Iscove’s Modified Dulbecco’s Medium
LRL: Lilly Research Laboratories
MBC: Metastatic breast cancer
NT: Not treated
PBS: Phosphate Buffered Saline
PI: Propidium Iodide
RT: room temperature
SD: Standard deviation
TNBC: Triple negative breast cancer
TR-FRET: Time-Resolved Foerster Resonance Energy Transfer
WO: Washout

## Acknowledgements

The authors wish to thank the donors, and the Biobank Hospital Universitario Puerta de Hierro Majadahonda (HUPHM)/Instituto de Investigación Sanitaria Puerta de Hierro-Segovia de Arana (IDIPHISA) (PT17/0015/0020 in the Spanish National Biobanks Network) for the human specimens used in this study. The authors thank Eglantine Julle-Daniere for her medical writing and editorial assistance for this manuscript, Dongling Fei for statistical analysis and data representation assistance, and Maria Jesús Ortiz for quality review assistance.

