## Supplemental Materials for "Efficient inhibition of cell proliferation and promotion of apoptosis requires continuous treatment with abemaciclib"

### Materials & Methods

#### Materials

Abcam: anti-CDK4 antibody (ab75511). Acros organics: PFA (119690010). Cell Signalling: lysis buffer (9803), and anti-CDK6 antibody (3136). Fisher: hiFBS (11082-147). Gibco: Ethylenediaminetetraacetic acid (EDTA, 15575-020), RPMI 1640 (A10491), hiFBS (16000-144), trypsin (25200-056), and PBS1X (70011-03). Invitrogen: IMDM (12440-053), RPMI 1640 (52400), and Pen/Strep (15140-130). Life Technologies: GMCSF (PHC2015); recombinant human full-length CDK4/Cyclin D1 protein, GST-tagged, expressed in insect cells (PR8064A, Lot# 880124H), tracer 236 (PV5592, Lot# 1351697 M), biotin-anti-His (PV6089, Lot# 1417453B), and Eu-Streptavidin (PV5900, Lot# 1610802F). Millipore: recombinant human full-length CDK6/Cyclin D3 protein (14-519, Lot# 2261733-A), and 96-well glass fiber filter plates (MAFCN0B50). Miltenyi: PI (130-092-052), Annexin V-FITC (130-092-052), Annexin buffer, Anti-H2AX pS139-FITC (130-107-584), Viobility 405/520 Fixable Dye (130-110-206), Inside Fix buffer (130-100-827), Permeabilization Buffer A (130-100-827), CD13-FITC (130-103-667), and CD11b-PE (130-091-240). Peprotech: IL3 (200-03). Perkin Elmer: 384-well Optiplate^TM^, MicroScintTM 20 (6013621), ULight-eIF4E-Binding Protein 1 (TRF0128-M, Lot# 1912472), and Eu-labeled anti-phospho-eIF4E-binding protein 1 (Thr37/46; TRF-0216-D, Lot# 1932859). Pierce: inhibitor cocktail (78446). ProQinase: CDK4/CycD1 (0142-0143-1). Sigma: Accutase (A6964), DMEM (D5796), Pen/Strep (P4458), Mytomycin C (10107409001), SCF (SRP3151), G-CSF (SRP3263), [gamma-33P] ATP (Adenosine 5’triphosphate) (NEN; 10µCi/µl, 3,000Ci/mmole) and ATP (Adenosine 5’triphosphate) (A-7699), MgCl_2_ (SM-2670), para-formaldehyde (F1635), ortho-Phosphoric acid (7664-38-2), Pen/Strep (P4458), DMEM (D5796), and tamoxifen (H7904). Upstate: C-terminal retinoblastoma fragment (CTRF; cat# 12-439). Deep well plates (278752) were purchased from Nunc, with 500 µl of cell medium per cell; six-well plates (RPMI 1640 A10491) were obtained from Gibco.

Compounds were prepared as 100% DMSO solution and diluted to generate 10-point 1:3 serial dilutions. DMSO solutions were diluted in kinase buffer (5 mM HEPES pH 7.5, 5 mM DTT, 0.01% TritonX-100 and 5 mM MgCl2) for biochemical assays as in the corresponding culture media for cellular experiments prior to be added to the enzyme reaction mix or cells.

#### Biochemical characterization

Kinetic (Ki) profiling - Ki determination

**Filter binding (FB) assays.** Compounds were mixed with substrate mix (C-terminal retinoblastoma fragment [CTRF] peptide and ATP/33-P ATP) at a final concentration range of 2µM-0.1nM. The mix was incubated for 90min at room temperature (RT) and the reaction was then stopped by adding 80 µl of 10% ortho-Phosphoric acid. The mix was next transferred to Multiscreen filter plates (Millipore) to retain phosphorylated peptide. The plates were read in a Microbeta Trilux instrument. Reaction mix with excess of EDTA was used as assay “bottom signal”; complete reaction mix without the inhibitor/compound was used as assay “top signal”.

For IC_50_ determination, results were fitted to a four-parameter dose-response curve using the following equation:

Y = bottom + [(top-bottom)/1+(x/IC_50_) ^slope^].

For the ATP saturation studies K_i_ and K’_i_ parameters were estimated using the formula:

v = V_max_[S]/[K_m_ (1+[I]/K_i_)+[S](1+[I]/K’_i_)]

#### Cell lines and culture condition

A panel of 32 BC cell lines (see Table S1 for details) was obtained from the American Type Culture Collection (ATCC, Manassas, Virginia, USA 30-4500 K) or the German Collection of Microorganisms and Cell Cultures GmbH (DSMZ). Cells were cultured according to ATCC or DSMZ recommendations for fewer than ten passages. Cells were seeded in 96 or 384-well plates and incubated overnight prior to treatment.

T47D cells (Figures 4, 5, S8 & S9) were obtained from ATTC (HTB-133) and were cultured in RPMI 1640 (Gibco), 10% hiFBS and 1% Pen/Strep (Sigma, P4458).

MDA-MB-453 cells (Figures S2&7) were obtained from ATTC® (HTB131™) and were cultured in DMEM (Sigma), 10% hiFBS (Gibco) and 1% Pen/Strep (P4458).

CD34+ cells (Figure 3, S4-6) were thawed and resuspended in IMDM, 10% hiFBS (Fisher), and 1% Pen/Strep (Invitrogen), supplemented with GMCSF 10ng/ml, G-CSF 10ng/ml, SCF 100ng/ml, IL3 10ng/ml and IL6 10ng/ml.

The IC_50_ of breast cancer cell lines was determine by curve fitting to a four-parameter logistic for each output using GENEDATA SCREENER® tool:

Y = bottom + [(top-bottom)/1+(x/ IC50) slope]

where Y = % inhibition (%Inh), X = concentration yielding y% inhibition, Bottom = minimum value of y attained by curve, Top = maximum value of y attained by curve, slope = steepness of curve at IC_50_. IC_50_: concentration of compound that reduces a given response (ligand binding, enzyme response) by 50%.

%Inh = [(median Max- x/ median Max – median Min)]/100

Resulting IC_50_ data for each treatment in each cell model were plotted as Geometric means in a waterfall plot using knime workflows. Biomarkers (ER or AR expression, HER2 amplification, or PIK3CA mutational) status was extracted from COSMIC db (COSMIC v79-Nov 2016).

#### In vitro drug treatment

Abemaciclib and its two main metabolites (internally synthesized) were added to the CD34+ progenitor cells at a concentration of 26 nM (Cmax, fu). After 13 days of maturation the cells per ml were measured using the flow cytometry technology. Data are normalized versus non-treated cells. The same procedure was also followed to investigate the effects of the combination of abemaciclib (100 nM) plus fulvestrant (30 nM) on the CD34 maturation process.

CD34+ bone marrow progenitor cells were treated with abemaciclib (250 nM), palbociclib (250 nM), or flavopiridol (250 nM) upon stimulation (IL3, GCSF, SCF, GM-CSF and IL6 cocktail), and using DMSO as non-treated controls. After 10 days of incubation cells were spun at 300 G, 5min and cell pellet was incubated with antibodies against CD13-FITC and CD11b-PE. After 10 minutes of incubation at RT protected from light, cells were washed with buffer (PBS, pH 7.2, 0.5% bovine serum albumin, 2 mM EDTA), spun at 300 G, 5 minutes, and supernatant was discarded. Cells were resuspended in buffer for acquisition in flow cytometer.

To measure the mitochondrial superoxide production after cell treatment, the Mitosox Red Mitochondrial Superoxide Indicator (Thermo, M36008) was used. For this, after cell detachment, the cells were allowed to recover in complete media at 37ºC for 20-30 mins. Then, cells were washed in PBS 1x and cell pellet was with 5 μM MitoSOX in HBSS/Ca2+/Mg2+, 10 minutes, 37ºC incubated following the vendors instructions. After the incubation time, cells were washed three times with HBSS/Ca2+/Mg2+ and prepared for acquisition in the cytometer (488-nm laser and 585/40 nm filter).

#### shRNA knockdown/phospho-RB Western.

MDA-MB-453 cells were seeded at a density of 400 000 cells per well in 6-well plates. CDK4 and CDK6 shRNA (Sigma, pLKO.1puro) lentivirus transduction was performed 24h post seeding with MOI of 10 in the presence of 10µg/ml polybrene. Cell media were changed to fresh complete medium after 24h transduction. Cells were collected 72h post transduction to test CDK4 and CDK6 knock down. 45µg of protein was loaded onto 5% Tris-Glycine gel for immunoblotting with anti-RB antibody (Cell Signaling 9309) and anti-phospho-RB antibody (pRB-S780 BD Pharmingen 558385), and 45µg of protein was loaded onto NuPAGE 4-12% Bis-Tris Gel for immunoblotting with anti-CDK4 antibody (Abcam ab75511) and anti-CDK6 antibody (Cell Signaling 3136) using ECL-HRP on Fujifilm LAS4000.

#### Analysis of cell viability

MDA-MB-453 CDK4 knockdown stable cell lines, generated with 1µg/ml puromycin selection, were plated in 96-well plates at 4 000 cells/well. On Day 1, 3, and 7, cells were fixed with fixative Prefer™ (Anatech, # 410) for 20min and then stained with 20µg/ml propidium iodide solution diluted in Phosphate Buffered Saline (PBS) containing 200μg/ml Ribonuclease A (Sigma R6513). The plates were scanned with ACUMEN EXPLORER™ to measure DNA content. To monitor senescence, cells were plated into 6-well plates at 150 000 cells/well. At Day 6, senescence was assessed using Cellular Senescence Assay Kit (Cell Biolabs, # CBA-230) per manufacturer’s instructions.

To measure apoptosis, CD34+ cells were washed with FACS Flow, spun at 300 G for 5min, and supernatant was removed. Cell pellet was incubated with Annexin V-FITC for 10min in dark following vendor’s instructions. Cells were then washed with Annexin buffer, centrifuged at 300 G for 5min, supernatant was discarded, and cell pellet was resuspended in 100 µl of Annexin Buffer before being analyzed in the cytometer. Cells were analyzed using flow cytometry (events were gated for debris exclusion and singlets selection). A minimum of 5,000 cells were analyzed per sample. PI (1:200) was added automatically by the cytometer. The percentage of cells at each apoptotic phase was represented: Annexin V-PI- (Alive). AnnexinV+/PI- (Early apoptosis), Annexin+/PI+ (Late apoptosis) and Annexin-/PI+ (Dead).

Anti-H2AX pS139-FITC expression was used to measure DNA damage. For this, after 13 days on treatment, cells were washed with PBS and cell pellet was incubated with Viobility 405/520 Fixable Dye for 15 min in the dark. Then, cells were washed and fixed using Inside Fix buffer for 10min at RT following vendor’s guidelines. Cells were then washed with PBS and cell pellet was permeabilized using Permeabilization Buffer A and incubated for 30min in ice. Next, cells were washed, and incubated with anti-H2AX pS139-FITC for 10min at RT. Finally, cells were washed and analyzed using the flow cytometer (gated on single cells and alive cells subpopulations). The mean fluorescence intensity, or median of fluorescence (MFI), was calculated, and compared for individual well and condition.

Cell cycle was analyzed using a Ki67/PI protocol. For this, after 13 days of treatment, cells were washed, fixed, and permeabilized following the same procedure as described above. Then, cells were labeled with Anti-Ki-67-Vio667 for 10min at RT. Next, cells were washed, and the cell pellet was incubated with PI/RNAse (Immunostep, PI/RNASE) for 15min at RT in the dark. Cells were analyzed using the flow cytometer (gated for debris exclusion and single cells). A minimum of 5 000 cells were analyzed per sample. The different phases of cell cycle were gated as follows: G0 (Ki67-/PIlow), G1 (KI67+/PIlow), S (Ki67+/PImed), G2M (Ki67+/PIhigh). All data were analyzed using FlowJo 10.6 and Graph Pad prism 8.

#### pRB at ser780 – High Content Imaging cell-based assay *[previously described in Torres-Guzmán et al. (2017)]*

MDA-MB-453 (CDK4-dependent) and NCI-H1568 (CDK6-dependent) cells were seeded at a density of 5 000 and 3 000 cells per well, respectively, in 96-well plates. Between 14-24h after seeding, cells were treated with compounds (abemaciclib, palbociclib, or ribociclib). Compounds were provided as 10 mM stock in 100% DMSO and were further diluted to achieve the desired final concentration (DMSO 0.2%).

After dosing with compounds, the cells were incubated at 37°C and 5% CO_2_ for 4 h before being washed once with PBS and adding 100 μl of fresh media. Cells were incubated once more for 1, 2, or 12 h at 37ºC/ 5% CO_2_ (washout [WO] step). After the WO, reference compound was re-added in minimal signal wells, to determine the minimal signal after WO (Figure S1).

To monitor the phosphorylation of Rb (ser780), a high content imaging cell-based assay was conducted, as previously used in Torres-Guzmán et al.[1]. After treatment, cells were fixed with 3.7% para-formaldehyde, permeabilized with cold Methanol and blocked with 1% BSA (Sigma) in PBS. Then, cells were treated with mouse anti-Rb (PS780) antibody (BD Pharmingen 558385) in 1% BSA in PBS overnight at 4°C. The next day, after several washing steps, cells were treated with goat anti-mouse IgG- Alexa Fluor™ 488 (Thermo Fisher A11008) in PBS for 1h at RT. After washing with PBS, 1:1,000 RNAase (Sigma R6513) and Propidium Iodide (PI) (Thermo Fisher P3566) dilution in PBS was added for 1h at RT. Fluorescence plates were scanned with ACUMEN EXPLORER™ monitoring Alexa Fluor 488 and PI signals using 488 nm wavelength.

#### Analysis and statistical considerations

Raw data were analyzed with FlowJo 10.6 software. Graph Pad v8.4.3 software was used for data analysis and representation of final readouts. JMP (Statistical Discovery from SAS) was used for the statistical treatment (ANOVA, pair-wise analysis) of the data.

**Flow cytometry analysis.** All analysis were carried out using the flow cytometry technology (Macsquant 10, Miltenyi):

- Cell number: cells were gated on size (FSC) and internal complexity (SSC) for debries exclusion and the cells per ml (gated on singlets) were used as final readout. The percentage of cell proliferation inhibition normalized versus non-treated cells and staurosporine maximum inhibition was represented.
- Apoptosis: Cells were washed with PBS 1X and the cell pellet was incubated with an antibody against Annexin V. After 10 minutes of incubation in dark, cells were washed twice using Annexin buffer V. Finally, cells were analyzed using the flow cytometer where PI (1:100) was automatically added. The percentage of cells at the different phases of apoptosis was represented (PI-/Annexin V- for Alive cells, PI-/Annexin V+ for Early apoptotic cells, PI+/Annexin V+ for Late apoptotic cells and PI+/Annexin V- for Dead cells).
- Senescence: For fluorescent detection of β-galactosidase, cells were washed with PBS 1X and fixed using 2% PFA (Acros organics, 119690010) for 10 minutes. Then, cells were washed using PBS+1%BSA and incubated with the cell even green reagent (Thermo, C10841) following the vendors indications (2h, 31ºC, no CO_2_). Finally, cells were washed and resuspended in 1% BSA in PBS for FACS analysis (488-nm laser and 525/50 nm filter). The results are represented as the percentage of green positive cells (gated on FSC, SSC, and singlets for debris exclusion and doublets respectively).

### Results

#### Potency of abemaciclib for CDK4: biochemicals and breast cancer cell-based assays

Abemaciclib is a more potent inhibitor of CDK4 than CDK6 in breast cancer cell lines

##### Biochemical characterization

In biochemical assay, abemaciclib inhibits the kinase activity of CDK4/cyclin D1 complexes with a Ki^ATP^= 0.6 nmol/l ± 0.3 nmol/l and of CDK6/cyclin D3 complexes with a Ki^ATP^= 8.2 nmol/l ± 1.1 nmol/l, showing that in an in vitro cell-free assay, abemaciclib demonstrates specificity of approximately 14-fold for CDK4/cyclin D1 over CDK6/cyclin D3 complexes (Table S1), regardless of the biochemical assay technic used (FB or TR-FRET). Under identical conditions, palbociclib did not exhibit such potency, and palbociclib could demonstrate a higher affinity to CDK6/cyclin D3 complexes as the IC_50_ for CDK6/cyclin D3 complexes was approximately 3-fold higher than the IC_50_ for CDK4/cyclin D1 complexes (Figure 1A).

**Supplemental Table S1**. **In vitro cell-free assays, abemaciclib shows different activity for CDK4 over CDK6 Biochemical data.** Head-to-head comparison of abemaciclib versus palbociclib and ribociclib. Kinetic parameters (Ki^ATP^) and selectivity ratio of abemaciclib for cyclinD1/CDK4 and cyclinD3/CDK6 complexes in vitro. Data reported as average of two independent determinations (n = 2) ± standard deviation (SD).

| **Method** | **Ki[ATP] FB (nM) ± SD** | | **IC_50_ FB (nM) ± SD** | | **Ki [ATP] TR-FRET (nM)** | |
| --- | --- | --- | --- | --- | --- | --- |
| **Compound** | **abemaciclib** | **palbociclib** | **abemaciclib** | **palbociclib** | **abemaciclib** | **palbociclib** |
| CDK4/CyclinD1 | 0.6 ± 0.3 ^(1)^ | 2.9 ± 2.6 | 1.9 ± 0.9 | 7.4 ± 0.3 | 0.7 | 9.2 |
| CDK6/cyclinD3 | 8.2 ± 1.1 ^(1)^ | 4 | 24.2 ± 10 | 1.9 ± 0.9 | 9.8 | 3 |
| **Ratio** | **14X** | **1.4X** | **13X** | **0.3X** | **14X** | **0.3X** |

(1) Torres-Guzmán R*.* et al. Oncotarget. 2017 Sep 19; 8 (41): 69493–69507

##### Cellular data

The effect of abemaciclib, palbociclib and ribociclib was further examined in two ER+ cell lines: one cell line CDK4-dependent (MDA-MB-453) and one cell line CDK6-dependent (NCI-H-1568). In the cellular assay, abemaciclib and ribociclib demonstrated a higher potency in the CDK4-dependent cell line, and abemaciclib was more potent than ribociclib in the CDK4-dependent cell line (Table S2). Consistent with the biochemical assay, palbociclib did not exhibit any selectivity towards CDK4/cyclin D1 complexes and demonstrated a similar level of inhibition in both cell lines (Figure 1C).

**Supplemental Table S2. Abemaciclib promotes more profound and durable effects in breast cancer cell lines.** IC_50_ values (nM) corresponding to the inhibition of pRb in breast cancer cells line indicated in the presence of abemaciclib, palbociclib, or ribociclib for 16 hours. Compounds were tested in duplicates within each individual experiment and tested on three separate occasions, except ribociclib that was tested (in duplicates) on two separate occasions

|  | **IC_50_ (nM)** | | |
| --- | --- | --- | --- |
| **Cell line** | **Abemaciclib** | **Palbociclib** | **Ribociclib** |
| **MDA-MB-231** | 17.11 ± 4.2 | 16.14 ± 7.6 | 71.53 ± 26.5 |
| **MCF-7** | 19.40 ± 7.4 | 25.56 ± 14 | 116.31 ± 78.5 |
| **BT474** | 4.58 ± 2.9 | 11.48 ± 5.5 | 33.99 ± 26.7 |
| **MDA-MB-175 VII** | 17.79 ± 2.8 | 18.91 ± 4.7 | 115.30 ± 80.9 |
| **EFM-19** | 4.40 ± 1.2 | 11.00 ± 0.6 | 31.01 ± 3.6 |
| **MDA-MB-361** | 8.24 ± 1.6 | 14.61 ± 2.1 | 54.25 ± 13.8 |
| **MDA-MB-134** | 8.50 ± 0.9 | 20.48 ± 8.2 | 59.96 ± 19.7 |
| **HCC70** | > 6000 | > 6000 | > 6000 |

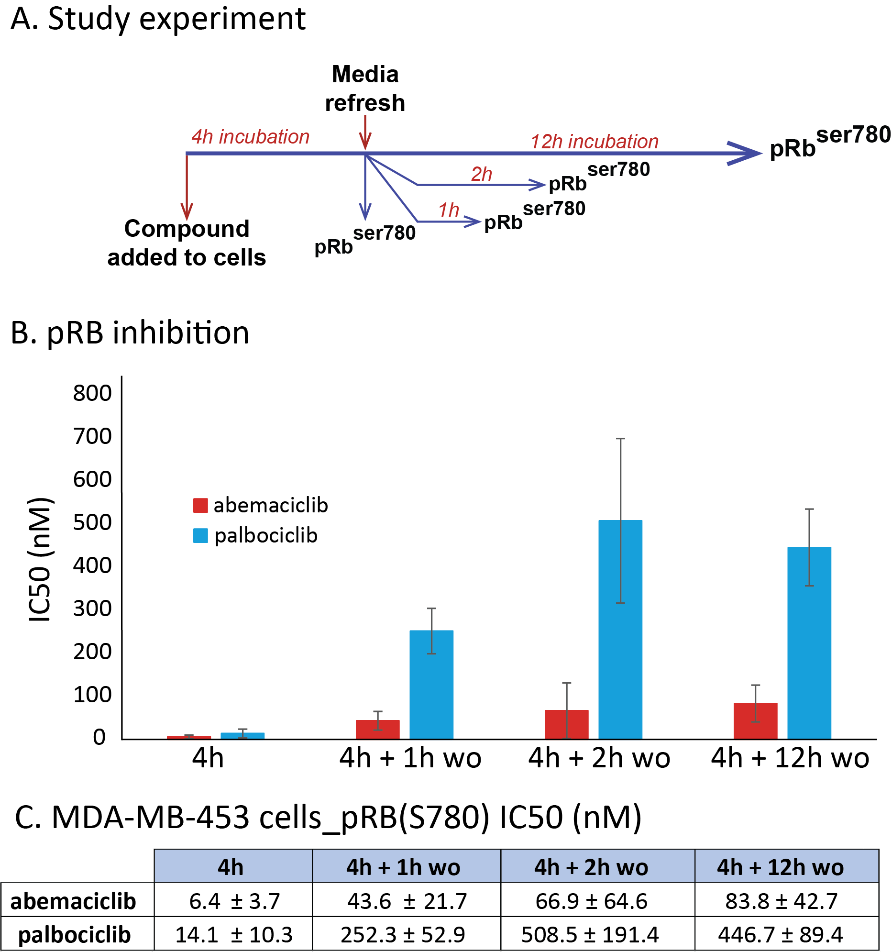

**Supplemental Figure S1.** **Abemaciclib promotes more profound and durable effects in breast cancer cell lines.** **(A)** Diagram of study experiment. IC_50_ values (nM) of washout experiment in MDA-MB-453 cells lines. Cells were treated for 4 hours with abemaciclib or palbociclib; **(B)** after compound removal, pRB inhibition was monitored at different time points (1, 2, or 12 hours). **(C)** Geometric means and SD for IC50 values are corresponding three separated experiments. pRb (S780): phospho-retinoblastoma at serine 780.

#### CDK4 plays a critical role in breast cancer

The role of CDK4 in cell proliferation was assessed in CDK4 knockdown stable cell lines generated in the AR+ breast cancer cell MDA-MB-453 (Figure S2A) and cell proliferation and senescence were assessed. The kinase activity of CKD4/cyclin D1 is involved in the cell cycle and triggers the progression from G1 to S1, and cell proliferation. In CDK4 knockdown cell lines CDK4-1 shRNA and CDK4-4 shRNA, the cells stopped growing and multiplying, thus indicating the interruption of cell proliferation (Figure S2B). CDK4 knockdown also led to cell senescence (Figure S2C). Taken together, these results demonstrate the dependency of the cell cycle on CDK4 in breast cancer cells expressing higher levels of CDK4. CDK4 is thus a key target in breast cancer therapy.

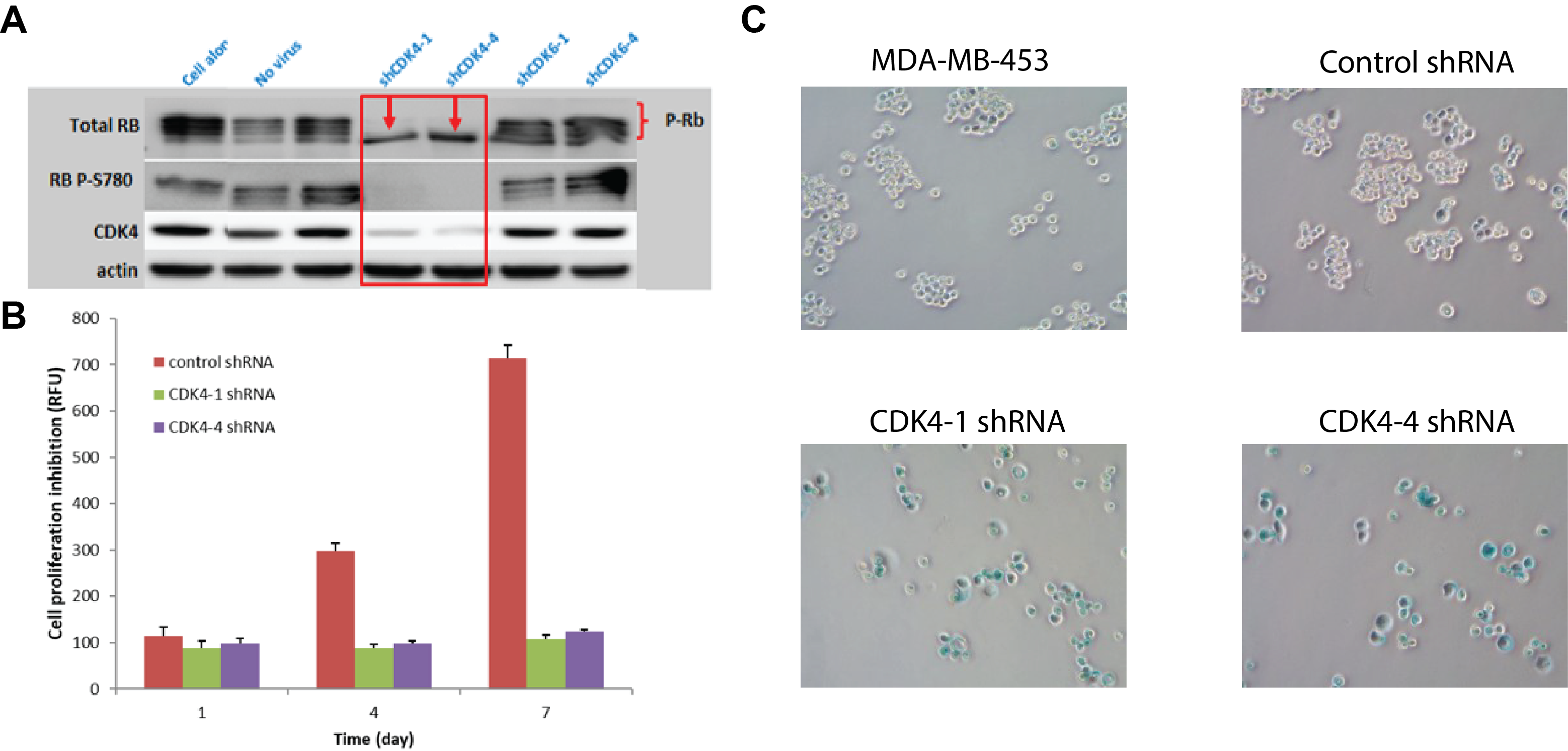

**Supplemental Figure S2.** **Knockdown experiment illustrating the role of CDK4 in breast cancer**. (**A**) shRNA Knock Down/Phospho-RB Western; (**B**) CDK4 knockdown inhibited cell proliferation in MDA-MB-453; (**C**) CDK4 knockdown induced senescence in MDA-MB-453 cells infected with shRNA lentivirus for 24; 48h after infection, cells were selected with puromycin for 6 days. Cells were replated in 96-well plate for proliferation assessment.

**Supplemental Table S3.** **Breast cancer cell lines.**

| **cell line** | **ER** | **HER2** | **TNBC** | ***BRAC1/2* ^(h)^** | ***RB1 mutations* ^(h)^** | **Rb expression ^(i)^** | ***PIK3CA mutations* ^(h)^** | **subtype** | **source** |
| --- | --- | --- | --- | --- | --- | --- | --- | --- | --- |
| BT-474 | ER+ | HER2+ |  | MUT BRCA2 (substitution-nonsense) | WT | + | MUT PIK3CA (substitution-nonsense) | luminal B | a,b,h, i |
| HCC1419 | ER+ | HER2+ |  | WT | WT | + | WT | luminal B | b, h, i |
| MDA-MB-361 | ER+ | HER2+ |  | MUT BRCA2 (substitution-missense) | WT | + | MUT PIK3CA (substitution-nonsense) | luminal B | b,h, i |
| UACC-3133 | ER+ | HER2+ |  | NA | NA | + | NA | luminal B | h, i |
| ZR-75-30 | ER+ | HER2+ |  | MUT BRCA2 (substitution-missense) | WT | + | WT | luminal B | b,h, i |
| CAMA-1 | ER+ | HER2- |  | WT | WT | + | WT | luminal B | b,h, i |
| EFM-19 | ER+ | HER2- |  | WT | WT | + | MUT PIK3CA (substitution-nonsense) | Luminal A | b,h, i |
| MCF-7 | ER+ | HER2- |  | NA | NA | + | MUT PIK3CA (substitution-nonsense) | Luminal A | a,h, i |
| MDA-MB-134-VI | ER+ | HER2- |  | NA | NA | + | WT | Luminal A | b,h, i |
| MDA-MB-175-VII | ER+ | HER2- |  | WT | WT | + | WT | Luminal A | b,h, i |
| MDA-MB-415 | ER+ | HER2- |  | WT | WT | + | WT | Luminal A | b,h, i |
| T-47D | ER+ | HER2- |  | NA | NA | + | MUT PIK3CA (substitution-nonsense) | Luminal A | b,h, i |
| ZR-75-1 | ER+ | HER2- |  | NA | NA | + | WT | Luminal A | b,h, i |
| UACC-812 | ER- | HER2+ |  | WT | WT | + | WT | HER2 | c, h i |
| AU565 | ER- | HER2+ |  | WT | WT | + | WT | HER2 | h, i, l |
| HCC1569 | ER- | HER2+ |  | MUT BRCA2 (substitution-missense) | WT | + | MUT PIK3CA (substitution-nonsense) | HER2 /post-EMT | b, h, i |
| HCC1954 | ER- | HER2+ |  | MUT BRCA1 (frameshift deletion) | WT | + | MUT PIK3CA (substitution-nonsense) | HER2/Basal | b, h, i |
| HCC202 | ER- | HER2+ |  | WT | WT | + | MUT PIK3CA (substitution-nonsense) | HER2 | b, h, i |
| HCC2218 | ER- | HER2+ |  | WT | WT | + | WT | HER2 | h, i, k |
| MDA-MB-453 | ER- | HER2+ |  | WT | WT | + | MUT PIK3CA (substitution-nonsense) | HER2 | a, e, h, i |
| SK-BR-3 | ER- | HER2+ |  | NA | NA | + | WT | HER2 | a, b, e, h, i |
| UACC-3199 | ER- | HER2+ |  | NA | NA | **-** | NA | HER2 | h |
| UACC-893 | ER- | HER2+ |  | WT | WT | + | MUT PIK3CA (substitution-nonsense) | HER2 | b, h, i |
| BT-20 | ER- | HER2- | TNBC | MUT BRCA2 (substitution-missense) | MUT (substitution missense) | + | MUT PIK3CA (substitution-nonsense) | basal | b, h, i |
| BT-549 | ER- | HER2- | TNBC | WT | WT | - | MUT PIK3CA (substitution-nonsense) | basal Claudin-Low | e, h, i |
| DU-4475 | ER- | HER2- | TNBC | WT | WT | - | WT | luminal | f, h, i |
| HCC1143 | ER- | HER2- | TNBC | WT | WT | + | WT | basal | b, h, i |
| HCC1187 | ER- | HER2- | TNBC | WT | WT | - | WT | basal | b, h, i |
| HCC1395 | ER- | HER2- | TNBC | MUT BRCA2 (substitution-nonsense) | WT | + | WT | basal | g, h, i |
| HCC1806 | ER- | HER2- | TNBC | WT | WT | + | WT | basal | b, h, i |
| HCC1937 | ER- | HER2- | TNBC | WT | MUT | + | WT | basal | b, j, g, h |
| HCC38 | ER- | HER2- | TNBC | WT | WT | + | MUT PIK3CA (substitution-nonsense) | basal | b, h, i |
| HCC70 | ER- | HER2- | TNBC | WT | MUT (deletion in frame) | - | WT | basal | b, h, i |
| Hs-578-T | ER- | HER2- | TNBC | WT | WT | + | WT | basal Claudin-Low/ post-EMT | a,b,h, i |
| MDA-MB-157 | ER- | HER2- | TNBC | WT | WT | + | WT | post-EMT | b, h, i |
| MDA-MB-231 | ER- | HER2- | TNBC | WT | WT | + | WT | basal Claudin-Low/ post-EMT | a,b,h, i |
| UACC-2087 | ER- | HER2- | TNBC | NA | NA | + | NA |  | h, i |

^a^ Breast Cancer: Basic and Clinical Research 2010:4 35–41; ^b^ Breast Cancer Res Treat (2007) 105:319–326; ^c^ Breast Cancer Res. 2011; 13 (6): R121; ^d^ Mol Cancer Ther. 2018 May;17 (5):897-907; ^e^ Holliday, D.L., Speirs, V. Choosing the right cell line for breast cancer research. Breast Cancer Res 13, 215 (2011); ^f^ J Cancer. 2017; 8 (16): 3131–3141; ^g^ Breast Dis. 2010; 32 (1-2): 35–48; ^h^ COSMIC.db (Nucleic Acids Research, Volume 47, Issue D1, 08 January 2019, Pages D941–D947); ^i^ internal data (WB); ^j^ Mol Cancer Ther; 16 (12) December 2017; ^k^ Breast Cancer Res 19, 65 (2017); ^l^ Am J Cancer Res. 2016; 6 (11): 2661–2678

ER: estrogen receptor; HER: human epithelial receptor; TNBC: triple-negative breast cancer

Signature for Rb expression in WB: (+) Visible band of Rb total and phosphoRb (ser780); (-) Total Rb band is not visible or barely visible with no sign of Rb phosphorylation pRb (ser780)

#### Breast cancer panel sensitivity to CDK4 & 6 inhibitors and biomarker analysis

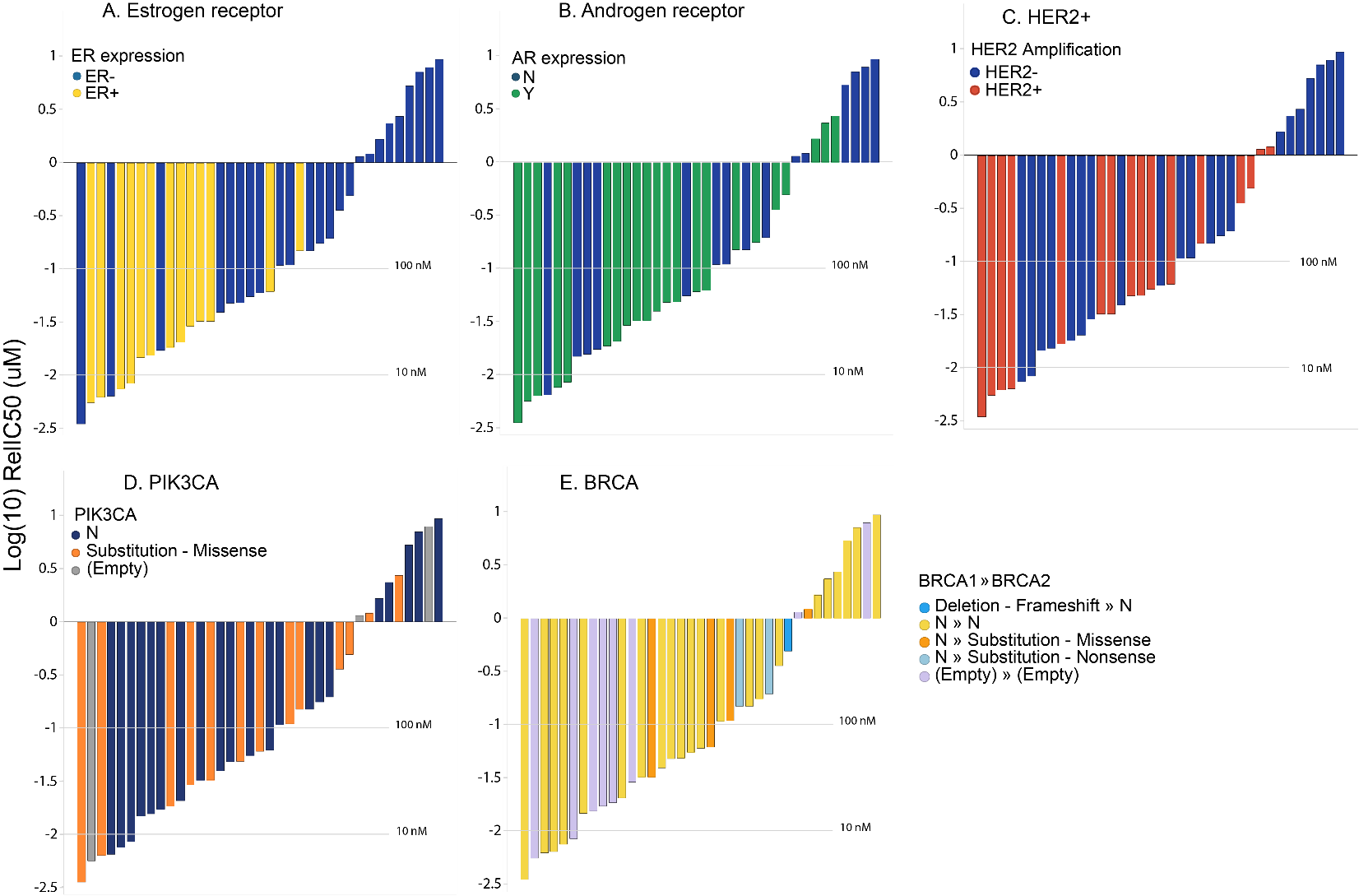
Abemaciclib inhibits cell proliferation in a wide range of breast cancer cell lines, showing activity regardless of HER2, PI3KCA and BRCA gene mutation status.

**Supplemental Figure S3.** Breast cancer cells sensitivity waterfall plots for abemaciclib related to (A) Estrogen receptor expression, (B) Androgen receptor expression, (C) HER2 expression, (D) PI3KCA and € BRCA gene activating mutations show sensitivity to abemaciclib. Biomarkers (ER or AR expression, HER2 amplification, or PIK3CA/BRCA mutational) status was extracted from COSMIC db (COSMIC v79-Nov 2016). Waterfall plots were generated using the geometric mean for each cell line and treatment.

#### Neutrophils maturation and neutropenia

CDK4 & 6 inhibitors share a common mechanism of action to induce neutropenia, which differs from the mechanism action of chemotherapy agents and others pan-CDKs inhibitors (paclitaxel or flavopiridol) (Figure S4). Abemaciclib metabolites, M2, and M20, show a similar effect on myeloid maturation of progenitor bone marrow cells. Cell surface markers (CD13 and CD11b) of myeloid maturation are less impacted by abemaciclib treatment than with palbociclib or a pan CDK inhibitor (Figure S5). The combination with fulvestrant does not increase the impact on in vitro hematopoiesis (Figure S6).

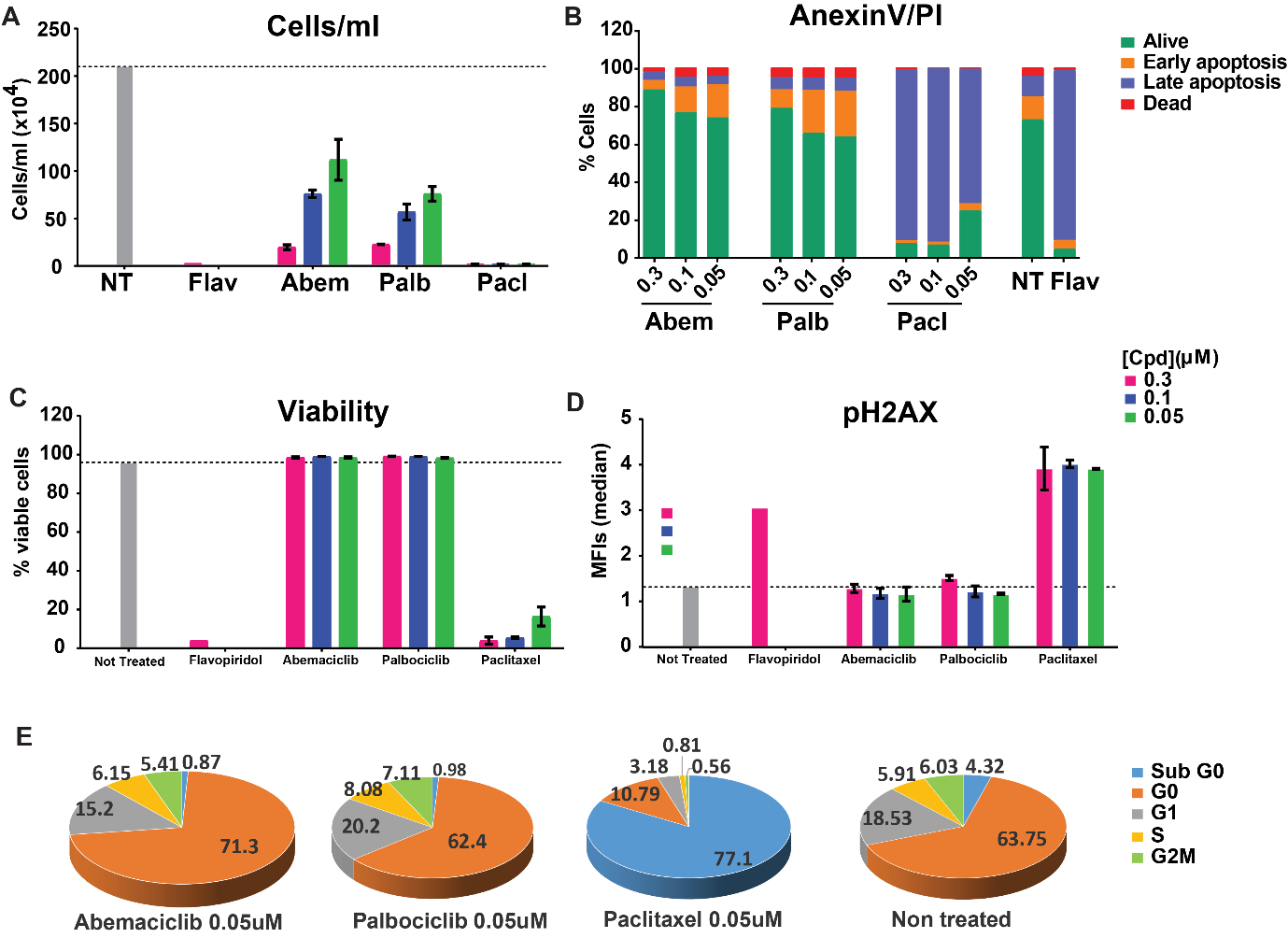

**Supplemental Figure S4.** **Different mechanism of action of abemaciclib, palbociclib, and paclitaxel on human bone marrow progenitor cells**. (**A**) Using flow cytometry cells/ml were counted after 13 days of CD34+ maturation in the presence of compounds. (**B**) Apoptosis of CD34+ cells was measured by running an Anexin V/PI assay. (**C**) Cell viability (vio fix dye 405/452) and (**D**) DNA damage (pH2AX) were also evaluated. (**E**) Cell cycle was measured using a Ki67/PI assay. The different CDK4/6 inhibitors share a common mechanism of action to induce neutropenia, which is distinct from that of chemotherapy agents or others pan-CDKs inhibitors.

NT: not treated; Abem.: abemaciclib; Flav.: flavopirodol; Palb.: palbociclib; Pacl.: paclitaxel

Flow cytometry analysis of stimulated healthy CD34+ cells (IL3, GCSF, SCF, GM-CSF and IL6 cocktail) for 10 days for myeloid cells to mature into neutrophils. Healthy CD34+ cells (Figure S5A) were treated with abemaclib (b), palbociclib (c) or flavopiridol (d) upon stimulation and using DMSO as not-treated controls (a). CD13 and CD11b surface markers were monitored as a reference for mature neutrophil. Phase I corresponds to myeloblast population, Phase II corresponds to promyelocytes population, Phase III corresponds to myelocytes and Phase IV corresponds to metamyelocytes and band neutrophils.

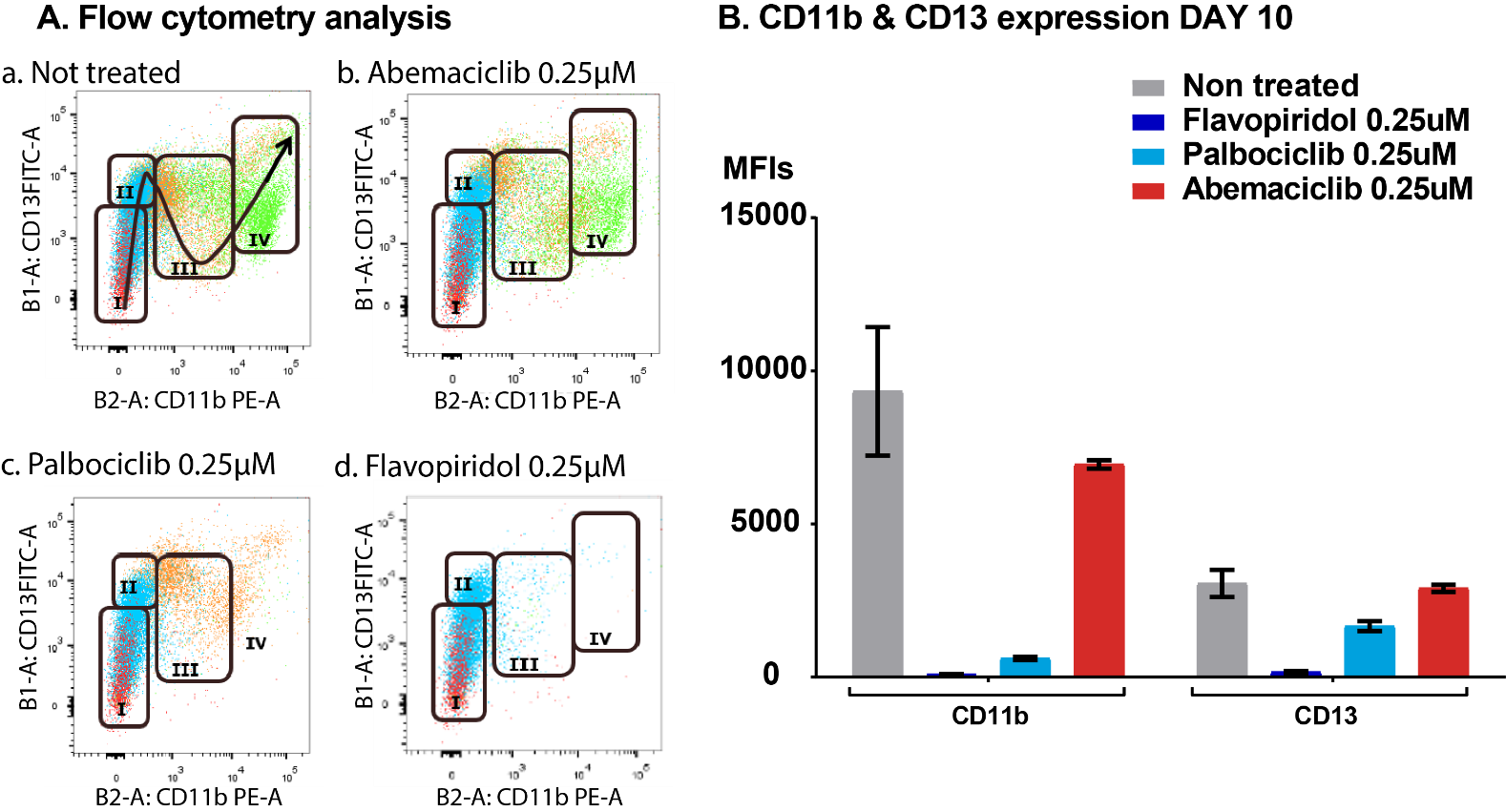

**Supplemental Figure S5. Impact of CDK4/6 inhibitors on neutrophils cell surface maturation markers. (A)** In the scatter plots the segregation of the different subpopulations can be observed under non-treated conditions (a), with 0.25uM abemaciclib (b), 0.250uM Palbociclib (c) or 250uM flavopiridol (d) as a known agent to promote neutropenia in-vivo by preventing the maturation of neutrophils. **(B)** The MFI for CD13 and CD11b biomarkers on Day 10 under different treatments is represented, which represents effects of abemaciclib, palbociclib, and flavopiridol in the final mature neutrophil outcome.

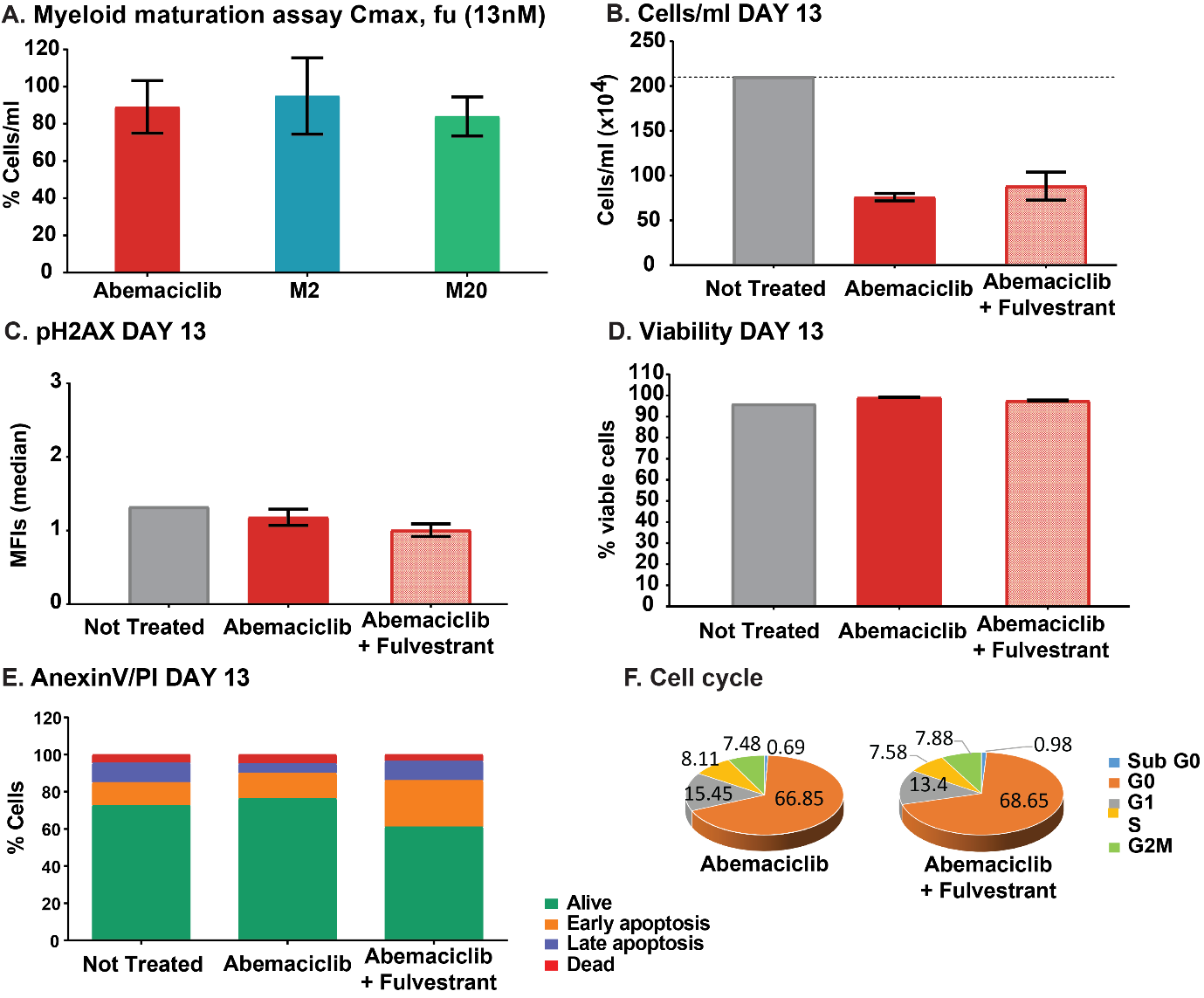

**Supplemental Figure S6.** **Metabolites and combo abemaciclib + fulvestrant impact on neutrophils.** (A) The two main metabolites of abemaciclib (M2 & M20) show a similar profile on myeloid maturation at the Cmax, fu (13 nM). (B-F) The combination of abemaciclib (100 nM) with Fulvestrant (30 nM) does not impact significantly the number of matured progenitor cells, neither the cell cycle, DNA damage, or viability. Only a small increase in early apoptotic cells was observed with the combination treatment.

#### Permanent exposure leads to durable effects after compound removal

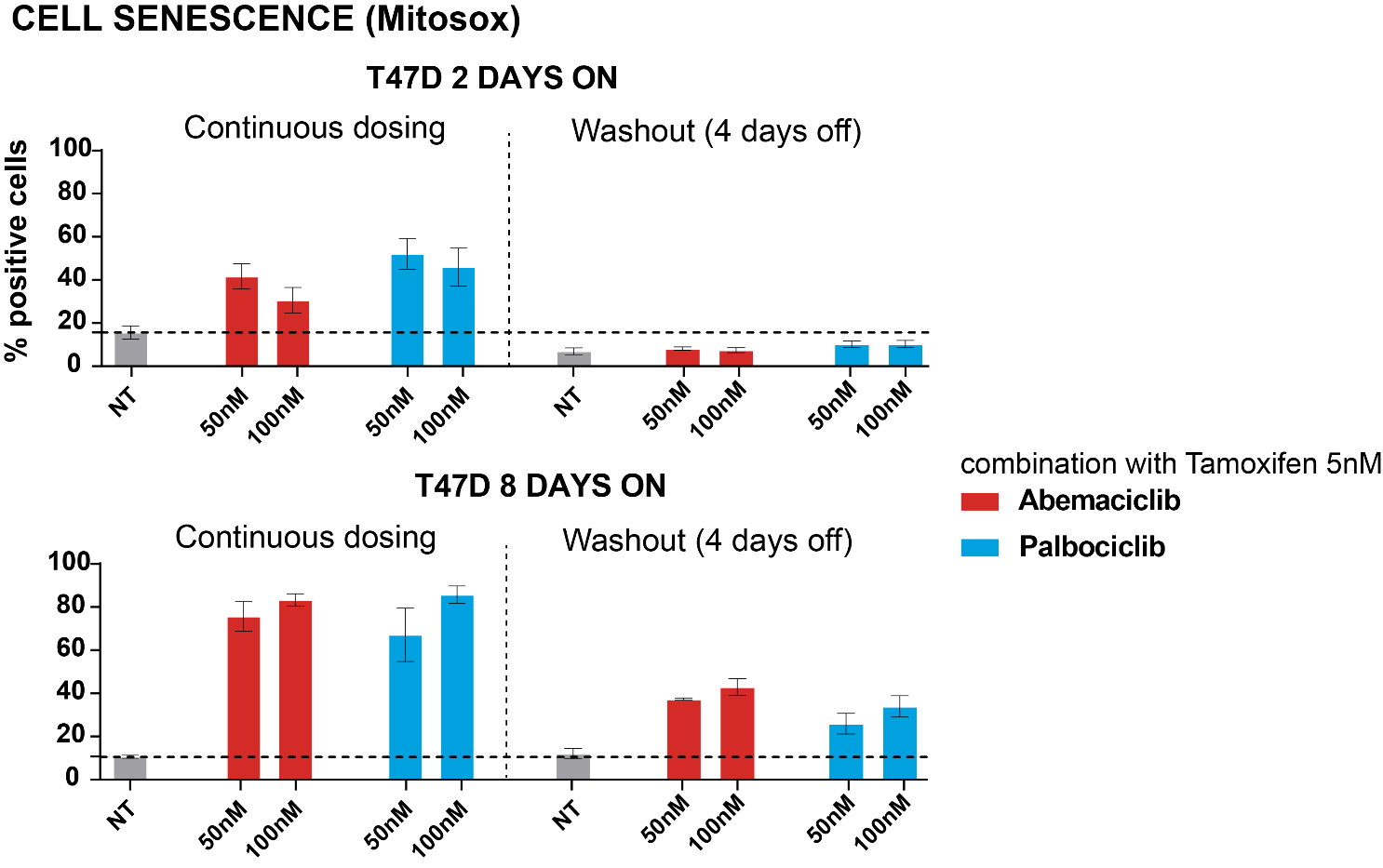

**Supplemental Figure S7.** **Washout studies demonstrated durable effects after compounds removal when cells are treated for a longer time. (A)** Assay design; T47D cells were treated with a combination of abemaciblib or palbociclib in combination with tamoxifen or fulvestrant for 2 or 8 days; compounds were removed, and readouts monitored after a 4-day washout period. **(B)** Percentages of senescent cells after combination treatment, monitored by Mitosox kit using flow cytometry.

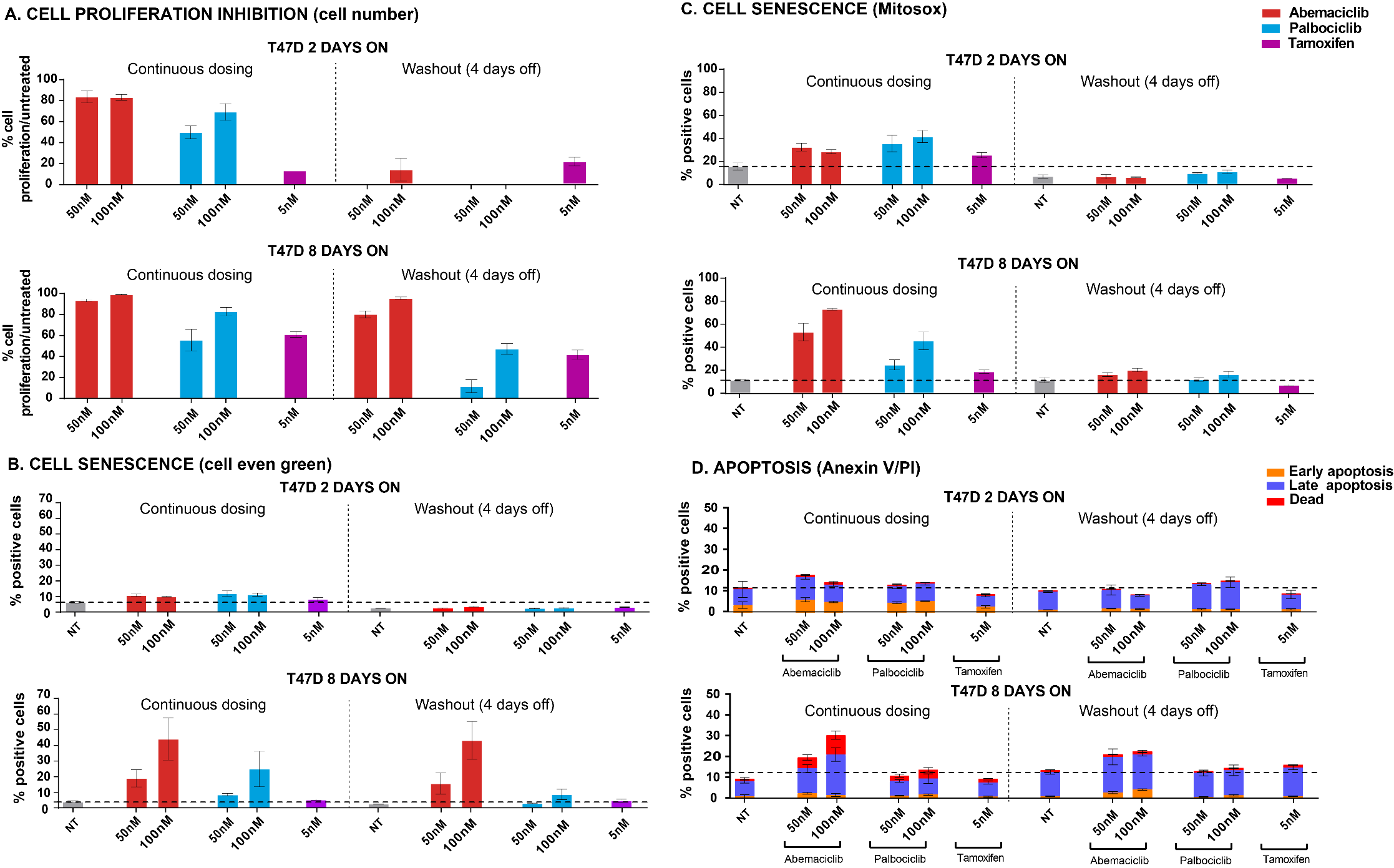
**Supplemental Figure S8. Abemaciclib showed a persistent effect when dosed as monotherapy for 4 days after compound removal. (A)** Cell proliferation inhibition measured as cells/ml by flow cytometry. **(B-C)** Percentages of senescent cells after single compound treatment, monitored by cell even green kit (B) or Mitosox kit (C). **(D)** Percentage of apoptotic cells after single compound treatment, monitored by Anexin V, and PI. The data are plotted as the mean (+/- SD) of three experiments.

### Table of Abbreviations

ATCC	American Type Culture Collection

BC	Breast cancer

BCA	Bicinchoninic acid

CTRF	C-terminal retinoblastoma fragment

DMSO	Dimethyl sulfoxide

DSMZ	German Collection of Microorganisms and Cell Cultures GmbH

EDTA	Ethylenediaminetetraacetic acid

ER	Estrogen receptor

ET	Endocrine therapy

FDA	Food and Drug Administration

HR	Hormone receptor

IMDM	Iscove's Modified Dulbecco's Medium

LRL	Lilly Research Laboratories

MBC	Metastatic breast cancer

PBS	Phosphate Buffered Saline

PI	Propidium Iodide

RT	room temperature

SD	Standard deviation

TNBC	Triple negative breast cancer

WO	Washout
